# Neonicotinoids and *Varroa* mites force the extending of longevity in a bee colony during overwintering to forget

**DOI:** 10.1101/2023.10.26.564284

**Authors:** Toshiro Yamada, Yasuhiro Yamada

**Affiliations:** Graduate School of Natural Science & Technology, Kanazawa University, Kakuma, Kanazawa, Ishikawa 920-1192, Japan; Department of Applied Physics, The University of Tokyo, 7-3-1 Hongo, Bunkyo-ku, Tokyo 113-0033, Japan

**Keywords:** seasonal changes, longevity, bee colony, overwintering, sugar-syrup pollen-paste, neonicotinoid, *Varroa* mite

## Abstract

A mathematical model that can estimate the apparent longevity of bee-colony proposed by the authors to clarify that the longevity normally changes with seasons as follows: The longevity maintains almost constant from at 20 to 30 days from late spring to late autumn, thereafter, it continues to extend till late spring up to 160 to 200 days. Immediately after overwintering, the longevity is shortened almost vertically from one sixth to one tenth. Such normal seasonal changes in longevity are shown when a pesticide-free food is administered to the bee-colony and when pesticide-containing sugar syrup is. However, abnormal seasonal changes in longevity, which does not extend even if winter approaches, are shown for the bee-colony ingesting neonicotinoid-containing pollen and for the colony infested with *Varroa* mites.

Judging from the fact that pollen is the main food for the bee-brood, that mites parasitize larvae and pupae, and that the vital functions and organs of honeybees are created during the larval and pupal stages, it can be inferred that a neonicotinoid-containing pollen paste and parasitic mites cause serious damage to the bee ability to detect the arrival of winter. Such dysfunction in the larval and pupal stages probably interferes with extending the longevity of adult bees even as winter approaches.

## Introduction

Lifespan of animal individuals is one of the important indicators for evaluating the activity of animals. In eusocial insects that form colonies such as honeybees, colonies can be treated as super-individuals that behave as if they were a single animal. It can be said that the day age (longevity) of the longest-living group of honeybees in a honeybee-colony is representative of the period of activity of the honeybee-colony. In other words, the activity period of a honeybee-colony is considered to be assessed by longevity which is an indicator of the life of the entire colony, rather than the lifespan of an individual bee.

The authors proposed a mathematical model (Y. Yamada et al., 2019) that estimates the longevity of the honeybee-colony only from the number of adult bees and the number of capped brood, and named the longevity estimated by the mathematical model “apparent longevity” in order to distinguish it from that measured directly. Using this proposed mathematical model and the number of adult bees and the number of capped brood measured in long-term field-experiments, we estimated seasonal changes of apparent longevity (Y. Yamada et al., 2019).

It was found that the apparent longevity of control (pesticide-free) colony estimated using long-term field-experiments in mid-west Japan extended rapidly during wintering, and that just before the end of wintering, the apparent longevity reached 6 to 10 times longer than that in non-winter seasons (Y. Yamada et al., 2019). Such a sudden change in apparent longevity cannot be explained only by the labor loads (Fukuda &Sekiguchi, 1966) such as foraging tasks (Schmid-Hempel &Wolf, 1988; Wolf & Schmid-Hempel, 1989) and nursing tasks (Amdam et al., 2009) that have been said so far. Therefore, the authors have proposed a hypothesis that a biological rhythm control system which is genetically pre-incorporated in a honeybee detects seasonal changes in the environment such as temperature, humidity and sunshine duration and results in causing seasonal changes in longevity (Y. Yamada et al., 2019).

In addition, the apparent longevity estimated from long-term field-experiment results on Maui without cold winters was quite different from the estimated results in mid-west Japan, and significantly fluctuated, but no seasonal changes were observed. In less seasonal-change areas such as Maui, the apparent longevity has been found to be greatly affected by physiological phenomena in honeybees, such as the absence of queens and a decrease in egg production (T. Yamada & K. Yamada, 2020).

There have been many reports on factors affecting the longevity of honeybees. Neonicotinoid pesticides (neonicotinoids) that appeared at the end of the 20th century are known to adversely affect the longevity of honeybees due to their characteristics (long-persistent, highly-insecticidal, systemic, neurotoxic). For example, neonicotinoids themselves directly shorten the longevity of honeybees (Tarek et al., 2018; Anderson et al., 2019; Aljedani, 2017), or indirectly shorten their longevity as follows:

Neonicotinoids can adversely affect the immunity of queen bees and cause the re-development of the disease (Brandt et al., 2017); they disrupt the circadian rhythms and sleep of honeybees and can impair honeybee navigation, time memory and social communication (Tackenberg et al., 2020); acetamiprid adversely affects memory-related properties in honeybees, homing ability and expression levels of memory-related genes (Shi et al., 2019). Neonicotinoids reduce the mating frequency of queens, resulting in a decrease in the genetic diversity of working bees, which can adversely affect colony vitality (Forfert et al., 2017); imidacloprid damages the development of the nervous system in areas responsible for both the sense of smell and vision in the larval stage of honeybees, inhibiting the activity of honeybees (Peng & Yang, 2016).

Moreover, *Varroa* mites also shorten the longevity of honeybees as follows: *Varroa* mites and deformed wing virus (DWV) reduce the lifespan of winter honeybees (Dainat et al., 2012); adult bees emerging from capped brood in the presence of *Varroa* mites cause their deformed wing and significantly reduce their longevity (Martin, 2013); external parasitic mites, *Tropilaelaps mercedesae*, reduce the longevity and emergence weight of honeybees (Khongphinitbunjong, 2016); *Varroa* destructor parasitic disease and DWV infections cause deformities during the development of Africanized bees, inhibiting cellular immunity and significantly reducing the lifespan of adult bees (Reyes-Quintana et al., 2019).

In addition, combination of the chronic exposure to the systemic pesticide (neonicotinoids and fipronil) and the infestation of the *Varroa* mite (Straub et al., 2019) or the microsporidian parasite Nosema ceranae which is deeply involved with the *Varroa* mite (Alaux et al. 2010; Vidau et al, 2011; Aufauvre et al., 2012; Aufauvre et al., 2014; Gregore et al. 2016) has synergistic and adverse impact on honeybee survival, and results in shortening the longevity of honeybee-colony.

Thus, while it has been found that the long-persistent and systemic pesticide (*e.g.* neonicotinoids) and the external parasitic mite (*e.g. Varroa* mites) and/or viruses mediated by those mites reduce the longevity of honeybees, we cannot find papers regarding the impacts of systemic pesticides and/or *Varroa* mites on seasonal changes in honeybee longevity other than in our study (T. Yamada & K. Yamada, 2020).

Therefore, since it is important to know what gives to seasonal changes in the longevity of bee colonies in beekeeping, this paper clarifies the following points using a mathematical model (Y. Yamada et al., 2019) that estimates the longevity of honeybee *Apis mellifera* colonies from only the numbers of adult bees and capped brood obtained from long-term field-experiments (T. Yamada et al., 2018a; 2018b; 2018c) conducted in the mid-west of Japan: **1)**Under the *Varroa* mite-free circumstances, on the difference in the seasonal changes in apparent longevity among the three kind of colonies, without pesticides (control) (hereafter, mite-free-CR), with the neonicotinoids dinotefuran (hereafter, mite-free-DF/SS) and clothianidin via sugar syrup (hereafter, mite-free-CN/SS) and with the organophosphates fenitrothion and malathion via sugar syrup (hereafter, mite-free-FT/SS and mite-free-MT/SS): **2)**On the difference in the seasonal changes in apparent longevity between two kinds of vehicles, sugar syrup (hereafter, SS) and pollen paste (hereafter, PP), to administer a pesticide: **3)**On the difference in the seasonal changes in apparent longevity between the mite-free-CR colony and the colony where no pesticide is administered and mites are infested (hereafter, mite-infested CR colony).

## Materials and Methods

For details, please refer to Methods S1-S3 in the supplementary materials.

### *Determination method of apparent longevity* (refer to Supplementary Method S1)

#### Mathematical Model

The authors have proposed a mathematical model that can estimate the apparent longevity from the numbers of adult bees and capped brood (Y. Yamada et al., 2019). Here a dynamic parameter called “apparent longevity”, *L (t)*, defined in Equation (S1) is introduced, which depicts the characteristics of the “boundary” day-age where the average honeybee-worker disappears from the bee colony:

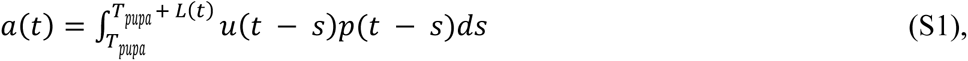

where *a(t)* and *u(t)* are the number of adult bees and the number of newly capped brood per unit time *t*, respectively. *p (t)* is the eclosion rate from capped brood (pupae in comb-cells with a sealed entrance) at time *t*. *T_pupa_* is the average period of capped brood, and the period of honeybees *Apis mellifera* is 12 days. If the bee colony is in a steady state, the worker bees die at the same age. *L(t)* represents the longevity of an adult bee. The age distribution of dying bees is given by the narrow Gaussian distribution.

Therefore, it is worth noting that Equation (S1) may provide an estimate of the longevity of a worker bee in a normal bee group.

In order to find the apparent longevity, *L(t)*, in Equation (S1), it is necessary to know *a(t)*, *u(t)* and *p(t)*. The values of *a(t)* and *u(t)* will be obtained from measurements in field experiments. Since it is difficult to directly measure *u(t)*, it is estimated from Equation (S2) using the *b(t)* value measured by the field experiment.

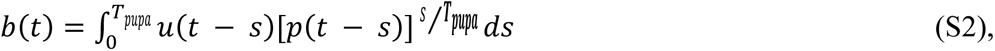

where *b (t)* is the number of capped brood at time *t*. Equation (S2) assumes that the pupa is removed as soon as it is dead in the comb cells.

By combining Equations (S1) and (S2), it is possible to characterize the honeybee group using a macroscopic amount on honeybees (the number of adult bees and number of capped brood) that are easy to get by experiments. Although *a(t)* in Equation (S1) and *b(t)* in Equation (S2) are continuous in the formula (it is a continuous function), it is actually impossible to measure them continuously in field experiments. Since the measurement date is discrete, Equations (S1) and (S2) were separated under the following assumptions.

1) The day-age distribution of capped brood on a measurement date is assumed to be discretely uniform during the capping period on the comb-cells (12 days), and to be constant between two consecutive measurement dates (in each measuring interval). These assumptions are equivalent to that the number of newly capped brood per day, *u(t)*, is constant between two consecutive measurement dates. If the number of capped brood measured on a measurement date is expressed in *b(t_k_)*, the daily average number of adult bees that newly emerges from the capped brood between one measurement date and the next measurement date is given by *b(t_k_)*/*T_pupa_*.

2) The eclosion rate is assumed to be constant in two consecutive measurement dates.

3) It is assumed that the pesticide intake per larva in two consecutive measurement dates is assumed to be the same as the intake of pesticides per bee. In our experiments, it is not possible to directly measure the intake of pesticides per larva between two consecutive measurement dates. Therefore, the pesticide intake per of the larva is substituted for that the that per measurable adult bee.

In Equations (S1) and (S2), *a(t)* and *b(t)* are known values measured in field experiments, but *L(t)*, *u(t)* and *p(t)* are unknown. In two simultaneous equations, one of these three unknown variables must be known before these simultaneous equations can be solved. Fortunately, the eclosion rates, *p(t)*, at various imidacloprid doses has already been reported (Yang et al. 2012). The eclosion rate required for calculating the apparent longevity can be obtained using the mean eclosion rate curve in Supplementary Figure S1.

Since the pesticide used in this experiment is dinotefuran, not imidacloprid, the dinotefuran concentration must be converted to the imidacloprid dose in order to use Supplementary Figure S1. Therefore, the imidacloprid dose equivalent to the insecticidal ability of a certain dinotefuran concentration is estimated as follows.

Field experiment conducted by the authors (T. Yamada et al., 2012) shows that the insecticidal ability of clothianidin for honeybees is about 2.5 times that of dinotefuran. It can be assumed that imidacloprid’s ability to exterminate honeybees is about 2.5 times that of dinotefuran, since the recommended insecticidal concentration of imidacloprid for exterminating stink bugs is approximately the same as that of clothianidin.

Pesticide intake per bee in two consecutive measurement dates can be estimated from the total pesticide intake and total number of adult bees between the measurement dates using the method proposed by the authors (T. Yamada et al., 2018a).

In the range of pesticide dosages in previous field experiments (see Supplementary Table S1), as shown in Supplementary Figure S2, the effect on the eclosion rate from bee pupae, *p(t)*, of the pesticide dose is small. That is, it can be assumed that the eclosion rate is equal and constant to 0.9 of the control group. Therefore, in this paper, the apparent longevity was calculated with a constant eclosion rate of 0.9 throughout the experiment as previously reported (T. Yamada & K. Yamada, 2020). Using the dataset at each measurement date obtained from the field experiment, the apparent longevity, *L(t)*, can be obtained by solving the discrete equation as shown in the previous report (Y. Yamada et al., 2019).

#### Determination Procedure of Apparent Longevity

The unknown variables in the equations for determining the apparent longevity of a bee colony in each measurement (observational experiment) are the apparent longevity *L(t)* and the number of the newly capped brood *u(t)*. Of these two variables, *u(t)* can be obtained from Equation (S2). Of these two variables, u(t) can be determined from Equation (S2).

Therefore, the *u(t)* value determined from Equation (S2) is substituted to Equation (S1), and the unknown variable *L(t)* in Equation (S1) is determined by an iteration method such as the Bisection method (Solanki et al., 2014) and the Simplex method (Nagata & Yamada, 1973). While changing the value *L(t)*, a certain value *L(t)* is repeatedly substituted for Equation (S1). And then, the calculations of discretization equations for Equations (S1) and (S2) are repeated until a difference between the calculated value of the number of adult bees a(t) obtained by the iteration method and the measured value of the number of adult bees by a field experiment reaches a given convergence criterion value (0.0001 in this paper) or less.

In the above calculation to seek the apparent longevity *L (t)*, actual measurement data can be obtained only for each measurement date. Therefore, the apparent longevity *L(t)* can be sought after the discretization of the equations while taking account of the life cycle from oviposition to eclosion of the honeybee (egg period 3 days; larval period 6 days; pupa (capped brood) period 12 days).

In this study, the apparent longevity *L(t)* is determined by the Bisection method (the convergence radius of L(t) is 0.0001 in this work). The calculations were performed using the Ruby programming language (Flanagan & Matsumoto, 2016). For more details such as the procedure and flowchart to seeking an apparent longevity, refer to Supplementary Method 1.

### Materials used in long-term field experiments

Neonicotinoid pesticides (dinotefuran, clothianidin) and organophosphate pesticides (fenitrothion, malathion) were used in long-term field experiments (T. Yamada et al., 2012; 2018a; 2018b; 2018c; 2018d; T. Yamada, 2020) as follows: Dinotefuran (neonicotinoid pesticide) (DF): Starkle Mate® (10% DF; Mitsui Chemicals Aglo, Inc., Tokyo, Japan). Clothianidin (neonicotinoid pesticide) (CN): DANTOTSU® (16% clothianidin; Sumitomo Co. Ltd., Tokyo, Japan). Fenitrothion (organophosphate pesticide) (FT): SUMITHION® emulsion (50% fenitrothion; Sumitomo Co. Ltd., Japan) was used. Malathion (organophosphate pesticide) (MT): MALATHON® emulsion (50% malathion; Sumitomo Co. Ltd., Japan).

Sugar syrup (SS) and pollen paste (PP) are used not only as food for bees, are often used but also as a vehicle for experiments in which pesticides are administered to bee colonies.

In this study, SS consists of 60 % sugar by weight and 40% water by weight, and was made by dissolving sugar in water at 70 °C. Sugar is granulated sugar that was composed of 99.7988% purified sugar (granulated sugar), Sodium chloride (salt) 0.1% or more, L-lysine hydrochloride 0.1% or more, and food dye (Blue No. 2) 0.0012% or more), and was purchased from the Japan Beekeeping Association (http://www.beekeeping.or.jp/). PP consists of 60% pollen by weight and 40% SS by weight, and was kneaded with a high-viscosity stirring bladed drill driver to a homogeneous paste at low speed while adding pollen to the SS.

For SS containing a pesticide, the desired amount of the pesticide was mixed into pesticide-free SS so that the desired concentration was achieved. PP containing a pesticide was made by kneading pollen into SS containing the desired pesticide concentration.

More details are described in Supplementary Method S2.

### Preparation for field experiment

In field experiments to investigate the effects of pesticides on bee colonies, it is necessary to minimize the effects of pesticides other than the pesticides administered in the experiments. Therefore, we tried to make the area around the experimental site as pesticide-free as possible. In other words, in the long-term field experiments in the Midwest of Japan, a water-feeding spot and foraging ground for bees were built in the immediate vicinity of the experimental site. Specifically, a system was installed that automatically supplies fresh water from a large tank. And, in a pesticide-free environment, pesticide-free plants and fruit-bearing trees were planted and cultivated. More details are described in Supplementary Method S2.

### Procedure for field experiment

The most important data in the field experiment of honeybee colony are both the number of adult bees and capped brood, which are indicators of the size, strength, vitality and power of the colony. To obtain these data as accurately as possible, the field experiment was carried out. The detailed procedure is described in Supplementary Method S2.

### Counting methods of adult bees, capped brood, mite-damaged bees

Using the photographs taken in field experiments, it was possible to determine the numbers of adult bees, capped brood and mite-damaged bees. Counting such numbers from photographs is an extremely difficult task. Therefore, with the cooperation of Mr. Yoshiki Nagai of Nanosystem Co., Ltd., Kyoto, Japan (http://nanosystem.jp/firm.htm), which develops image processing software, the author has developed computer software in 2012 that automatically counts the relevant numbers from photographic images, and improvements have been made to the software since. The accuracy and operation of the counting have been greatly improved. By using this software, the time and accuracy for counting such numbers have been significantly improved, compared to the amount of work required to count by hand directly from the original photo.

However, these counts are still inaccurate because of bee overlap, out-of-focus images, extreme contrast differences in the images, misjudgments of the counting targets, and so on. In this software, it is also possible to manually correct after automatic counting. Therefore, after automatic counting, the counting mistakes due to the automation were corrected by manual operation while zooming into the image, and the corrected number became the final data. Thus, errors still emerge in automation when using this software, but by using manual operation, it is possible to correct the counting errors. The improvements in the counting accuracy and speed under this software contributed greatly to the improvement of the data analysis accuracy and the speed in the field experiments.

When counting the number of *Varroa* mites, it is impossible to count *Varroa* mites that causes damage to a brood in a comb cell. If any damage is received from *Varroa* mites during the brood stage, there should be a trace of the damage even if the brood becomes an adult bee. Therefore, assuming that such traces confirm the presence of mites, we checked the adult bees for traces of mites. Based on the criteria for determining the traces due to *Varroa* mites (Supplementary Figure S5), the number of the traces (including *Varroa* mite itself) is counted on the photographic image while being enlarged with the help of the automatic counting system newly developed. More details on counting methods of the numbers of adult bees, capped brood and mite-damaged bees are described in Supplementary Method S3.

## Results

Here, we will examine the following seasonal changes in apparent longevity of colonies, which was estimated though a mathematical model (Y. Yamada et al., 2019) using the numbers of adult bees and capped brood measured in long-term field experiments (T. Yamada et al., 2018a; 2018b; 2018c; 2020).

### Mite-free-CR colony

The mite-free-CR colony (2011/2012 CR-1; 2012/2013 CR-1; 2012/2013 CR-2; 2013/2014 CR-1; 2013/2014 CR-2) (T. Yamada et al., 2018a; 2018b; 22018c) shows seasonal changes in apparent longevity that begins to extend around late September, becomes the longest (about 160 to 200 days) just before the end of overwintering, becomes instantly short up to near low levels (20 to 30 days) just after overwintering, and then stays almost constant at low levels, albeit with small fluctuations (Figure 1A; Supplementary Figures S7 to S9) (Y. Yamada et al., 2019). Such seasonal changes in apparent longevity are normal in areas with four seasons until the colony extinction.

**Figure 1.**
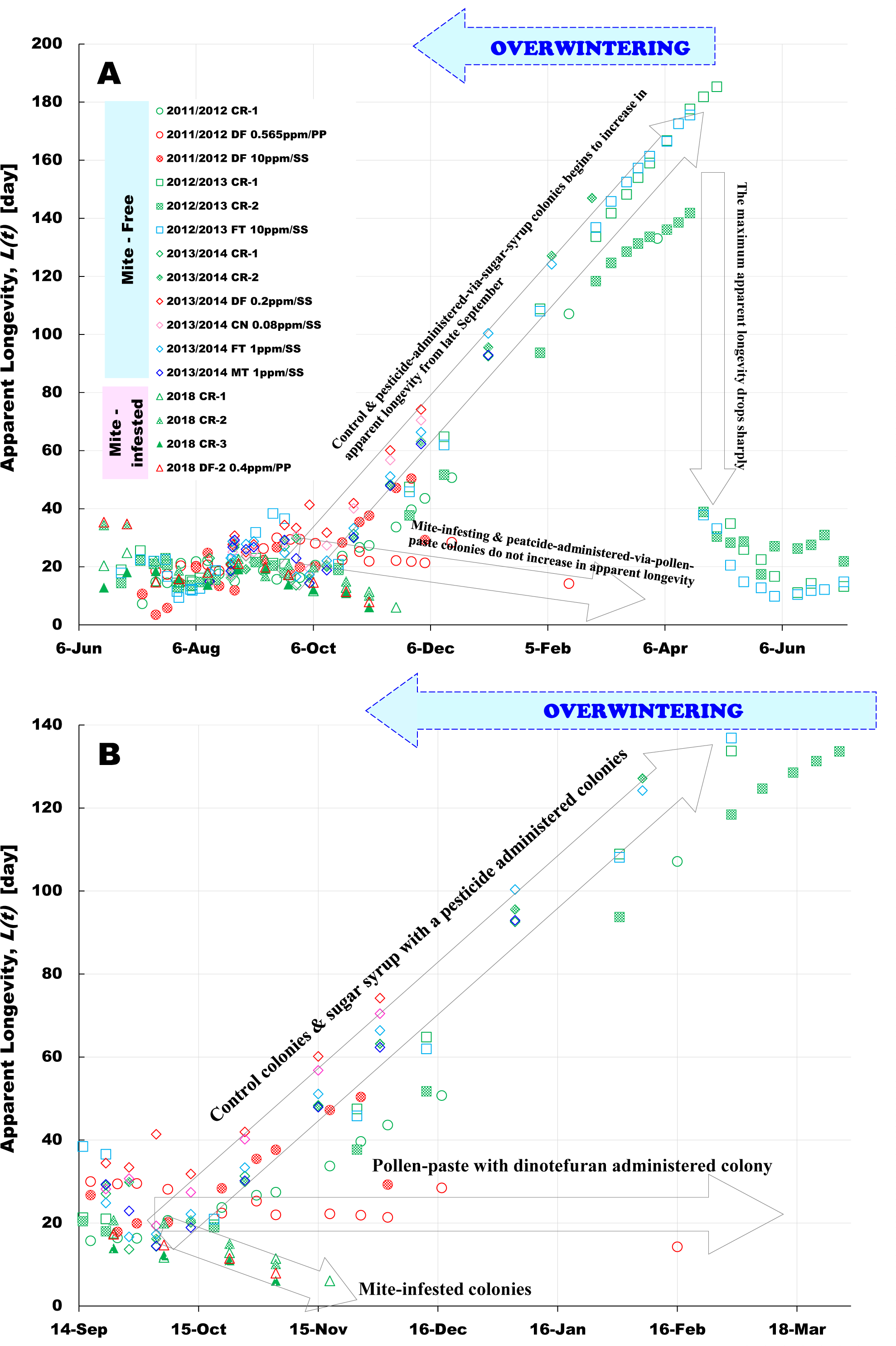
seasonal changes in apparent longevities of honeybee colonies in four long-term field-experiments. (**A**): General view. (**B**): Date-axis enlarged view. Apparent longevities were estimated by using the numbers of adult bees and capped brood obtained from four long-term field-experiments conducted in : 2011 to 2012 (T. Yamada et al., 2018a) for the colonies of “2011/2012 CR-1”, “2011/2012 DF 0.565ppm/PP” and “2011/2012 DF 10ppm/SS”; 2012 to 2013 (T. Yamada et al., 2018b) for the colonies of “2012/2013 CR-1”, “2012/2013 CR-2” and “2012/2013 FT 10ppm/SS”; 2013 to 2014 (T. Yamada et al., 2018c) for the colonies of “2013/2014 CR-1”, “2013/2014 CR-2”, “2013/2014 DF 0.2ppm/SS”, “2013/2014 CN 0.08ppm/SS”, “2013/2014 FT 1ppm/SS” and “2013/2014 MT 1ppm/SS”; 2018 (T. Yamada, 2020) for the colonies of “2018 CR-1”, “2018 CR-2”, “2018 CR-3” and “2018 DF-2 0.2ppm/PP”. The colonies which had become extinct before wintering (before late November) are eliminated in this study. Where CR, DF, CN, FT and MT denote control (pesticide-free), dinotefuran, clothianidin, fenitrothion and malathion, respectively. “201x/201y” denotes the experimental period from 201x to 201y, “X Yppm” denotes the concentration “Y ppm” of the pesticide “X”. “syrup” and “pollen” denote SS and PP of vehicles through which the pesticide is administered into the honeybee-colony, respectively. See Supplementary Tables S3 to S6.

Such normal changes in apparent longevity that extends as winter approaches cannot be always explained by the conventional theories such as workload on honeybees (Schmid-Hempel &Wolf, 1988; Amdam et al., 2009; Omholt, 1988) and nutritional intake (Alqami, 2006; Wang et al., 2014; Yang et al., 2017); for example, the facts that the apparent longevity begins to rise around late September when honeybees are active; that the apparent longevity continues to extend during the overwintering season when bees don’t always have enough food and even when a queen bee begins to oviposit in mid-February, and so on. The extending of apparent longevity as winter approaches can be caused by the genes already embedded so as such facts are steered.

### Mite-free-neonicotinoids/SS (mite-free-DF/SS and mite-free-CN/SS)

The mite-free-DF/SS colony (2011/2012 DF 10ppm/SS; 2013/2014 DF 0.2ppm/SS) (T. Yamada et al., 2018a; 2018c) and the mite-free-CN/SS colony (2013/2014 CN 0.08ppm/SS) (T. Yamada et al., 2018c) show seasonal changes in apparent longevity that begins to extend around late September as in case of the mite-free-CR colony but decrease rapidly soon after the beginning of the overwintering because of the colony extinction (Figure 1A; Figures S7 & S9). The colony extinction can be inferred to be caused by the weakening of the colonies due to the long-persistent toxicity of dinotefuran or clothianidin.

### Mite-free-organophosphates/SS (mite-free-FT/SS and mite-free-MT/SS)

The mite-free-FT/SS colony (2012/2013 FT 10ppm/SS) (T. Yamada et al., 2018b) exhibits normal seasonal changes in apparent longevity and success in overwintering almost the same as in case of the mite-free-CR colony (Figure 1A; Supplementary Figure S8). On the other hand, The mite-free-FT/SS colony (2013/2014 FT 1ppm/SS) (T. Yamada et al., 2018c) and the mite-free-MT/SS colony (T. Yamada et al., 2018c) exhibit seasonal changes in apparent longevity that begins to extend around late September as in case of the mite-free-CR colony but reduce sharply in the middle of the overwintering because of the colony extinction (Figure 1A; Supplementary Figure S9). The colony extinction can be inferred to be caused by the slight-weakening of the colonies due to the short-term lasting toxicity of fenitrothion and malathion.

### Mite-free-DF/PP

The mite-free DF/PP colony (2011/2012 DF 0.565/ppm) (T. Yamada et al., 2018a) shows abnormal seasonal changes in apparent longevity that does not begin to extend even late September and keeps approximately constant till the colony extinct (Figure 1B; Figure S7). The colony extinction can be inferred to be caused by the weakening of the colonies due to the long-persistent toxicity of dinotefuran.

### Mite-infested-CR

The mite-infested-CR colony (2018 CR-1; 2018 CR-2; 2018 CR-3) (Yamada, 2020) also shows abnormal seasonal changes in apparent longevity. Far from beginning to begin to extend late September, the apparent longevity begins to decrease and the colonies become extinct after the start of overwintering (Figure 1B; Supplementary Figure S10).

### Mite-infested-DF/PP

The mite-infested-DF/PP colony (2018 DF-2) (2020) also shows abnormal seasonal changes in apparent longevity in the same as in case of the mite-infested-CR; far from beginning to begin to extend late September, the apparent longevity begins to decrease and the colonies become extinct after the start of overwintering (Figure 1B; Supplementary Figure S10).

## Discussion

Focusing on how abnormal changes in apparent longevity are caused, we will discuss various differences of seasonal changes in apparent longevity.

### The difference in seasonal changes in apparent longevity among mite-free-CR, mite-free-DF/SS and mite-free-CN/SS, mite-free-FT/SS and mite-free-MT/SS

Until the colony extinction, the apparent longevity of both the mite-free colony where no pesticide is administered and a pesticide is administered via SS shows almost the same normal-changes of the seasons. When the colony is strong, it can succeed in overwintering, and when the colony is weakened by the pesticide or others, it will fail in overwintering (Figure 1A; Supplementary Figures S7 to S9). The reason why the pesticide administered to the colony via SS cannot cause abnormal changes in apparent longevity will be probably because SS (honey) is an energy source. Incidentally, genes, biological functions and organs are created in egg, larval and pupal stages of honeybees. In addition, SS (honey) is ingested for bees’ activities mainly by adult bees. but not ingested almost by a queen bee, larvae and pupae. Therefore, SS can hardly cause abnormal changes in AL.

### The difference in seasonal changes in apparent longevity between two kinds of vehicles (SS, PP) to administer a pesticide under the mite-free circumstances

The mite-free colony where a pesticide is administered via SS (2011/2012 DF 10ppm/SS; 2013/2014 DF 0.2ppm/SS; 2012/2013 FT 10 ppm/SS; 2013/2014 1ppm/SS; 2013/2014 MT 1ppm/SS) exhibits normal seasonal changes in apparent longevity that begins to extend from late September and continues to extend till the colony becomes extinct during overwintering. On the other hand, the mite-free colony where a pesticide is administered via PP (2011/2012 DF 0.565ppm/PP) exhibits abnormal seasonal changes in apparent longevity; the apparent longevity keep approximately constant in all seasons, even during overwintering, till the colony extinct as if the colony cannot discern the approach of winter (Figure 1B; Supplementary Figures S7 to S9).

The difference in the vehicle to administer a pesticide can cause the above difference in seasonal changes in apparent longevity.

Honey (SS) is the staple food of adult bees and pollen (PP) is the staple food of larvae and pupae. In addition. royal jellies which are made from pollen are ingested by queen bees and young larvae. Eggs are oviposited by a queen bee and functions and organs of honeybees are made in larval and pupal stages. From the above, pesticides involved in pollen can adversely affect honeybees during gene transcription, functional differentiation and organ creation. Long-persistent pesticides with high toxicity and neurotoxicity, such as neonicotinoids, will continue to adversely affect honeybees over a long period of time once they are ingested during the egg, larval and pupal stages. Such long-term adverse impact can cause gene transcription errors or some kind of dysfunction such as biological rhythm disorder. Such impediments will make it difficult to predict the approach of winter.

### The difference in seasonal changes in apparent longevity between the mite-free-CR colony and the mite-infested CR colony

The mite-free-CR colony (2018 CR-1; 2018 CR-2; 2018 CR-3) exhibits normal seasonal changes in apparent longevity, while the mite-infested-CR colony exhibits abnormal seasonal changes in apparent longevity similar to the mite-free-DF/PP colony (2011/2012 DF0.565ppm/PP) (Figure 1B; Supplementary Figures S7 & S10).

It can be assumed that the difference between these seasonal changes in apparent longevity is due to the fact that *Varroa* mites attack larvae and pupae (Sammataro, et al., 2000), in whose stages functions and organs of honeybees are made, and cause damage them. It can be assumed that mites and vector-borne viruses in the larval and pupal stages disrupt or destroy the honeybee’s biological clock and sensory organs, with making it impossible to recognize seasonal changes.

Though seasonal changes in apparent longevity of mite-infested colonies should hold constant in all seasons till the colony extinction in this case, the fact that they decrease from mid-September till the colony extinction can be caused by the extreme weakening of the colonies due to higher than 50% of mite-prevalence (Figure 2). In addition, there is little difference in seasonal changes in apparent longevity between the mite-infested CR colony and the mite-infested DF/PP colony. The reason will be that the influence of damages by mites is too strong.

**Figure 2.**
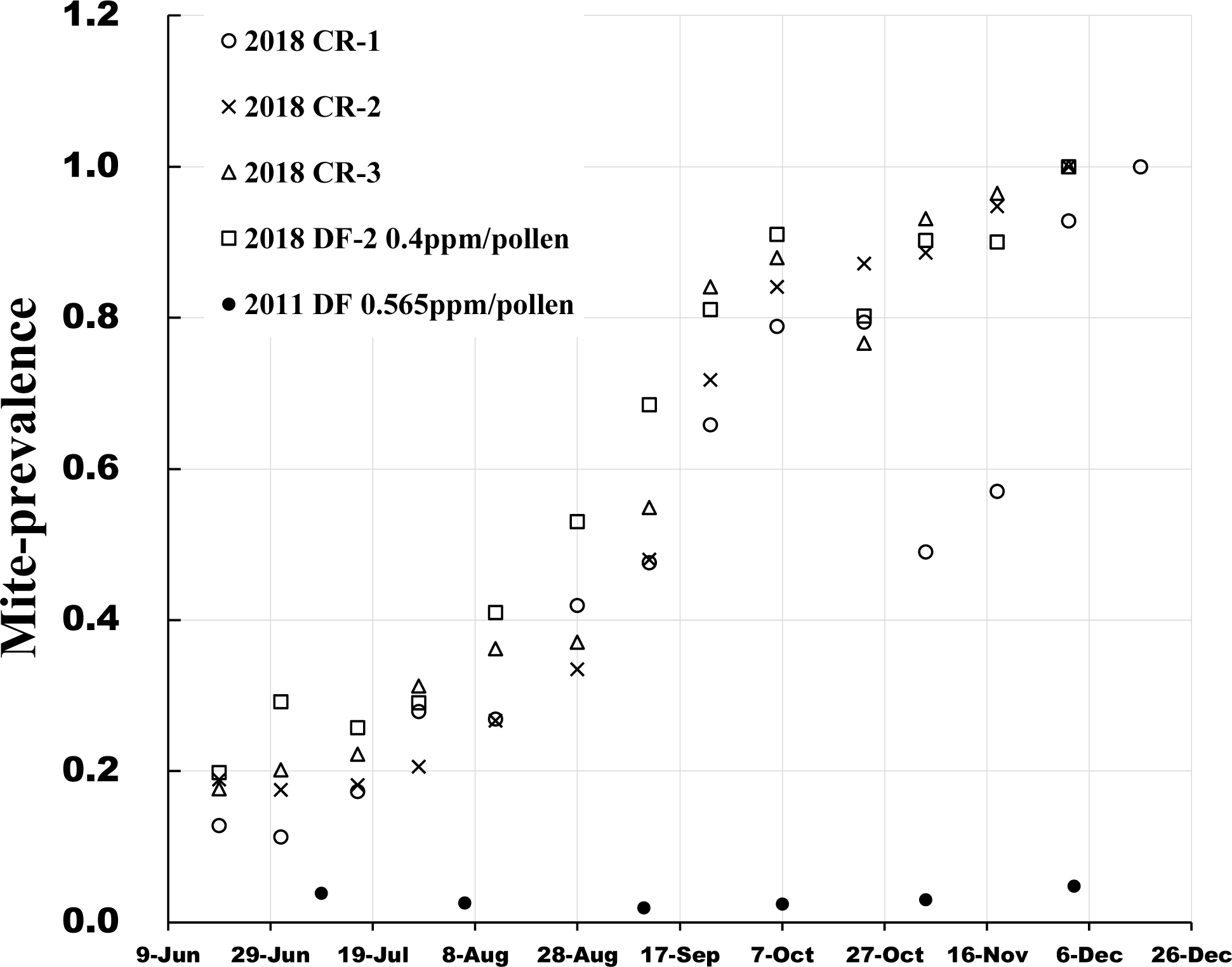
Comparison of mite-prevalence between the mite-infested colonies and the mite-free-&-toxic-PP colony. 2018 CR-1, 2018 CR-2 & 2018 CR-3: Pesticide-free control colonies in 2018-Experiment (CR-1, CrR-2, CR-3); 2018 DF-1, 2018 DF-2 & 2018 DF-3: Experimental colonies where dinotefuran is administered via PP in 2018-Experiment (DF-1, DF-2, DF-3). 2011/2012 DF: Experimental colonies where dinotefuran is administered via PP in 2011/2012-Experiment. Mite-prevalence denotes the ratio of bees damaged by mites to total bees in each measurement interval. The apparent longevity of all colonies in this figure could not begin to extend around late September. Mite prevalence of all colonies (2018 CR-1, 2018 CR-2, 2018 CR-3, 2018 DF-1, 2018 DF-2, 2018 DF-3) in 2018-Experiment, where dinotefuran (neonicotinoid) were administered via PP, are about 50% in mid-September, about 80% in mid-October, and finally more than 90% (called mite-infested colony). On the other hand, the mite-prevalence of experimental colony in 2012-Experiment (2011/2012 DF) where dinotefuran was administered via PP is almost 2∼3%, and at most less than 5%. See Supplementary Table S7.

Incidentally, it is suspected that the fact that seasonal changes in apparent longevity in mite-free DF/PP colony may be due to mite-infection. Therefore, to confirm the infection status of mites, the number of damaged bees by mites in the mite-free-DF/PP colony was counted according to the sections 1.3.3 and 1.3.4 in Supplementary Method S3. As a result, it turned out to be that the mite-free-DF/PP colony is hardly infested with mites because its mite-prevalence is 5% or less, while that of the mite-infested colony reaches over 90% (Figure 2). It was verified that the previous report (T. Yamada et al., 2018a) that there are no mites is correct, and the apparent longevity constant seasonal-change of the mite-free--DF/PP colony is not due to the infestation of mites, but to the ingestion of dinotefuran via PP.

## Conclusions

The mite-free CR colony and the colony where a pesticide is administered via SS colony exhibit normal seasonal changes in apparent longevity that begin to extend from late September till the colony extinction, while the mite-infested colony and the mite-free DF/PP colony exhibit abnormal seasonal changes in apparent longevity that keep constant in all seasons or decrease even when winter approaching till the colony extinct. Such abnormal seasonal changes in apparent longevity are inferred to be caused by serious damages on eggs, larvae and pupae.

## Supplementary material

Supplementary Methods S1 to S3; Supplementary Figures S1 to S10; Supplementary Tables S1 to S7.

## Acknowledgments

We would like to greatly appreciate the hearty cooperation of Mr. Yuhki Nagai (Nanosystem Co., Ltd., Kyoto, Japan) for developing and improving automatic counting software to improve the counting speed and accuracy of the numbers of adult bees, capped brood and damaged mites. We would like to thank for the cooperation of Mr. Kazuo Harada (Platon Co., Ltd., Kakogawa, Japan) when improving the input and output of software with apparent longevity. We would also like to express their deep gratitude to the following people for their devoted cooperation in conducting the field-experiments and counting the numbers of adult bees and capped brood whose measurements proved to be highly accurate through the estimation of apparent longevity with a mathematical model; Ms. Kazuko Yamada; Ms. Hiroko Nakamura; the members of YUINOTE (Mr. Tetsuya Kojima, Ms. Yuki Kojima, Dr. Hiroshi Mibayashi, Ms. Kimiko Mimura-Ruth, and Mr. Kenji Mimura) which is a natural farming group in Noto, Japan. We appreciate also Ms. Kazuko Yamada’s cooperation in algorithmic design for longevity estimation. Dr. Ken Hashimoto (Okayama, Japan) gave us his warm and appreciate advice, guidance and encouragement when we had some troubles on this research. Thank you deeply. Finally, we would like to thank everyone who gave us this opportunity and/or assisted us for our research.

## Supplementary Material

### 1 Materials and Methods

#### 1.1 Supplementary Method S1: Determination method of apparent longevity

##### 1.1.1 Mathematical model

Using the numbers of adult bees and capped brood obtained from long-term field experiments, the authors have proposed a mathematical model for estimating the apparent longevity of a honeybee colony at every observation-experiment interval. There are various day-age groups in the bee group, of which the theory seeks the day-age of the oldest living in honeybee group. The following is an overview of the mathematical model.

If we can estimate the apparent longevity of a honeybee colony, we can get more comprehensive information from field experiments. The authors have proposed a mathematical model that can estimate the apparent longevity from the numbers of adult bees and capped brood. Here a dynamic parameter called “apparent longevity”, *L (t)*, defined in Equation (S1) is introduced, which depicts the characteristics of the “boundary” day-age where the average honeybee-worker disappears from the bee colony:

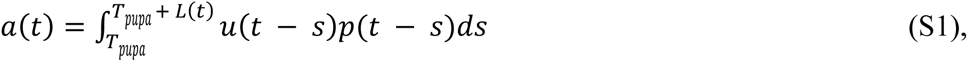

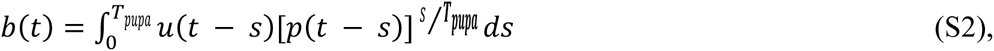

2) The eclosion rate is assumed to be constant in two consecutive measurement dates.

In Equations (S1) and (S2), *a(t)* and *b(t)* are known values measured in field experiments, but *L(t)*, *u(t)* and *p(t)* are unknown. In two simultaneous equations, one of these three unknown variables must be known before these simultaneous equations can be solved. Fortunately, the eclosion rates, *p(t)*, at various imidacloprid doses has already been reported (Yang et al. 2012). Figure S1 shows the relationship between the imidacloprid dose and the eclosion rate (mean, minimum, maximum value) in a semi-logarithmic graph. The eclosion rate required for calculating the apparent longevity can be obtained using the mean eclosion rate curve in Figure S1.

Since the pesticide used in this experiment is dinotefuran, not imidacloprid, the dinotefuran concentration must be converted to the imidacloprid dose in order to use Figure S1. Therefore, the imidacloprid dose equivalent to the insecticidal ability of a certain dinotefuran concentration is estimated as follows.

Field experiment conducted by the authors (T. Yamada et al., 2012) shows that the insecticidal ability of clothianidin for honeybees is about 2.5 times that of dinotefuran (this is the same as the ratio of the recommended insecticidal concentration of dinotefuran for exterminating stink bugs to that of clothianidin).

It can be assumed that imidacloprid’s ability to exterminate honeybees is about 2.5 times that of dinotefuran, since the recommended insecticidal concentration of imidacloprid for exterminating stink bugs is approximately the same as that of clothianidin. Pesticide intake per bee in two consecutive measurement dates can be estimated from the total pesticide intake and total number of adult bees between the measurement dates using the method proposed by the authors (T. Yamada et al., 2018a).

In the range of pesticide dosages in previous field experiments (see Table S1), as shown in Figure S2, the effect on the eclosion rate from bee pupae, *p(t)*, of the pesticide dose is small. In other words. there is little difference in calculated apparent longevity between when the eclosion rate changes with the pesticide dose per bee in the period of the interval between arbitrary two consecutive measurement dates as expressed by *p(t) = f (pesticide dose)* and when it is assumed that the eclosion rate is equal and constant to 0.9 of the control group.

Therefore, in this paper, the apparent longevity was calculated with a constant eclosion rate of 0.9 throughout the experiment as previously reported (T. Yamada & K. Yamada, 2020). Using the dataset at each measurement date obtained from the field experiment, the apparent longevity, *L(t)*, can be obtained by solving the discrete equation as shown in the previous report (Y. Yamada, 2019).

##### 1.1.2 Determination procedure

The procedure for calculating the apparent longevity using the proposed mathematical model as described above will be described below. The unknown variables in the equations for determining the apparent longevity of a bee colony in each measurement (observational experiment) are the apparent longevity *L(t)* and the number of the newly capped brood *u(t)*. Of these two variables, *u(t)* can be obtained from Equation (S2). Of these two variables, *u(t)* can be determined from Equation (S2).

Therefore, the *u(t)* value determined from Equation (S2) is substituted to Equation (S1), and the unknown variable *L(t)* in Equation (S1) is determined by an iteration method such as the Bisection method (Solanki et al., 2014), the Simplex method (Nagata & Yamada, 1973) and so on as below. While changing the value *L(t)*, a certain value *L(t)* is repeatedly substituted for Equation (S1). And then, the calculations of discretization equations for Equations (S1) and (S2) are repeated until a difference between the calculated value of the number of adult bees *a(t)* obtained by the iteration method and the measured value of the number of adult bees by a field experiment reaches a given convergence criterion value (0.0001 in this paper) or less.

In the above calculation to seek the apparent longevity *L(t)*, actual measurement data can be obtained only for each measurement date. Therefore, the apparent longevity *L(t)* can be sought after the discretization of the equations while taking account of the life cycle from oviposition to eclosion of the honeybee (egg period 3 days; larval period 6 days; pupa (capped brood) period 12 days).

In this study, the apparent longevity *L(t)* is determined by the Bisection method (the convergence radius of *L(t)* is 0.0001 in this work). The calculations were performed using the Ruby programming language (Flanagan & Matsumoto, 2016). The flowchart is shown in Figures S3 and S4. Figure S3 is a whole flow diagram for determining an apparent longevity *L(t)*, and Figure S4 shows a procedure for seeking the apparent longevity *L(t)* in Equation (S1) by the Bisection method. These flowcharts are described in a little more detail below.

Here, we will explain the whole flow of the calculation to seek the apparent longevity (Figure S3). The eclosion rate *p_k_* of a honeybee colony is calculated using the relationship between the dose of imidacloprid per larva and the eclosion rate (Figure S1) (Yang et al. 2012). Our previous experiments were conducted using pesticides other than imidacloprid. In addition, the intake of a pesticide by an adult bee can be directly calculated from experimental data, but that by alarva cannot. Larvae cannot directly take food (sugar solution, pollen paste) and adult bees (nurse bees) usually feed larvae on food. Therefore, it can be supposed that the intake of a pesticide of one larva is almost the same as that of an adult bee. In addition, it is assumed that the pesticide intake per bee is the same regardless of the role (nursing, forming) in adult bees. In other words, “dose of imidacloprid per larva” is equivalent to “intake of imidacloprid per bee”.

By the way, imidacloprid and other pesticides (dinotefuran, clothianidin, fenitrothion, malathion) have different insecticidal ability for bees, respectively. With considering the insecticidal ability of each pesticide, each pesticide intake is converted into an intake of imidacloprid intake, and the ratio of the dosage concentration of each pesticide to the bee colony is adjusted so that the insecticidal activity to exterminate stink bugs, which are common pests in Japan, is the same (T. Yamada & K. Yamada, 2020; T. Yamada et al, 2018b; 2018c).

In a honey bee colony during the period of each interval between two consecutive measurement dates, the total amount of a pesticide ingested by adult bees and the total number of adult bees involved in the pesticide intake are different at each interval. Incidentally, the total number of adult bees during the period of each interval can be obtained by the sum of the both numbers of adult bees that will newly eclose from capped brood and the number of adult bees existing from the beginning of each interval. Exceptionally, the total number of adult bees during the period of the final interval is obtained by the sum of the both numbers and the number of capped brood at the end of experiment. Because the capped brood can be presumed to have already ingested the pesticide.

Therefore, it is necessary to determine the intake of a pesticide per bee for each interval (see below for details). If the intake of a pesticide per bee at each interval is known, the eclosion rate *p_k_* in each interval can be estimated from Figure S1.

In a honeybee colony, by dividing the total amount of pesticide intake (*TI_k_*) ingested by the honeybee colony between the two consecutive measurement dates by the sum (*N_k_*) of the total number of adult bees (*B_k_*) eclosing between the (*k−1*)^th^ measurement date to the *k*^th^ one and the measured number of adult bees (*a_k−1_*) already existing on the (*k−1*)^th^ measurement date, the average intake of a pesticide per bee (*I_k_* = *TI_k_* / *N_k_*) can be determined. In each measurement interval, the intake of a pesticide per bee is converted to that intake of imidacloprid per larva under the assumptions mentioned above, and then the eclosion rate *p_k_* is estimated from Figure S1. In the pesticide-intake range in our field experiments, the apparent longevity *L(t)* calculated with the estimated eclosion rate *p_k_* depending on the pesticide intake differs little from the apparent longevity calculated with the constant eclosion rate of 0.9 (Figure S2) on the assumption that the eclosion rate of pesticide-intake colony is independent of pesticide intake and equal to the eclosion rate for the control colony. Therefore, the eclosion rate of 0.9 is used to calculate the apparent longevity in this paper.

Next, taking into account the life cycle of the bee (egg period of 3 days; larval period of 6 days; pupa (capped brood) period of 12 days), the simultaneous equations that discretize Equation (S2) on the *k^th^* measurement date is solved, and the number of newly capped brood *u_k_* is determined (**STEP 1**).

The number of newly capped brood *u_k_* determined as described above is substituted for Equation (S1). Then, while changing the apparent longevity *L_k_* on the *k^th^*measurement date with the Bisection method, calculations are repeated till both a difference between the measured number of adult bees and the calculated one and a difference between the old value of *L_k_* and the new one becomes both less than 0.0001. When these convergence criteria are satisfied, *L_k_* is the determined value. After iterating the above calculations and completing them for all the number of experiments (k= 1, 2, 3, ··, N) (**STEP 2**), the calculated results are printed.

When determining the apparent longevity *L_k_* by the Bisection method, first *L_k_* is increased with a rough step size (0.5 in this study), and two points (*L_k_* values) where the signs of the Y ((*a_k_*)_measured_ − (*a_k_*)_calculated_) are reversed are found (Figure S4). The two *L_k_* values before and after the sign is reversed are sandwiched, and the step size is further reduced, and the calculations are repeated until both the Y is closest to zero with the Bisection method and the absolute value of (*L_k, new_* - *L_k, old_*) becomes less than 0.0001. When the above criteria are satisfied, the value of *L_k_* is regarded as the determined apparent longevity.

#### 1.2 Supplementary Method S2: Field experiments

##### 1.2.1 Precautions for obtaining accurate measurements

Both of the numbers of adult bees and capped brood, which are very important for obtaining apparent longevity by use of a mathematical model and can be measured in the experiment, are deeply involved through the life cycle of honeybees each other. If the measurement accuracy of either number is poor, a contradiction will arise between these two measurements. For example, when some of adult bees have gone out foraging, the number of adult bees will underestimate the actual number existing in the honey bee colony. In this case, because there is an unreasonable difference in numbers between the two, apparent longevity cannot be accurately obtained when solving simultaneous equations in a mathematical model, or cannot be sometimes even determined because the simultaneous equations cannot be solved. Therefore, experiments are conducted while paying attention to the following points to measure accurately the numbers: The experiment is started immediately after dawn to minimize the number of adult bees out of the hive-box. The comb-frame with bees is pulled out from the hive-box as gently as possible, when taking picture, to prevent the bees from flying away, and is gently set on the photo stand. The experiment is conducted while avoiding a windy or rainy day, judging from the weather forecast, because of preventing the bees from becoming restless. In more details, see the previous paper (T. Yamada, 2020).

##### 1.2.2 Materials used in the field experiments and their preparation

The following materials were used in the long-term field experiments.

###### Pesticides

1) Dinotefuran (neonicotinoid pesticide) (DF): Starkle Mate® (10% DF; Mitsui Chemicals Aglo, Inc., Tokyo, Japan).

2) Clothianidin (neonicotinoid pesticide) (CN): DANTOTSU^®^ (16% clothianidin; Sumitomo Co. Ltd., Tokyo, Japan).

3) Fenitrothion (organophosphate pesticide) (FT): SUMITHION® emulsion (50% fenitrothion; Sumitomo Co. Ltd., Japan) was used.

4) Malathion (organophosphate pesticide) (MT): MALATHON^®^ emulsion (50% malathion; Sumitomo Co. Ltd., Japan).

###### Foods which are sometimes used as vehicles to administer a pesticide to a bee colony

1) Sugar syrup (SS): Granulated sugar was purchased from the Japan Beekeeping Association (http://www.beekeeping.or.jp/) and was composed of 99.7988% purified sugar (granulated sugar), Sodium chloride (salt) 0.1% or more, L-lysine hydrochloride 0.1% or more, and food dye (Blue No. 2) 0.0012% or more). A total of 20 kg of granulated sugar and 13.33 kg of hot water at about 75 ℃ were mixed in a 50 L plastic tank, and then 60% SS of granulated sugar was produced.

2) Pollen paste (PP): In total, 25 kg of pollen imported from Spain was purchased from Tawara Apiaries Co., Ltd., Kobe, Japan (https://tawara88.com/about.html). The pollen was used after it was lightly ground with “Kona Ace A-7” flour milling equipment manufactured by Kokkousha Co., Ltd., Nagoya, Japan (http://www.kokkousha.co.jp/). The viscosity of PP was determined by the ratio of SS in PP, the particle size of the pollen, and the temperature of the PP. The ratio of the pollen and SS that did not fall when the PP-filled tray was turned upside down was preliminarily examined. As a result, 60% pollen by weight and 40% SS by weight was confirmed to be an appropriate ratio. Pesticide-free PP and PP containing pesticide of desired concentration were prepared in 10 kg each. PP (6 kg of pollen and 4 kg of SS) was prepared in a large plastic bucket by kneading with a drill screwdriver with stirring blades used for high viscosity at a low speed until achieving a uniform paste.

3) SS with pesticide of a desired concentration: It was produced using 60% SS solution containing the desired concentration of pesticide and sugar. The prepared SS containing pesticide in a 10 L-container was blocked from light using a black bag and stored in a refrigerator.

4) Preparation and maintenance of the tray filled with PP: The 300 g PP, after being weighed in an upper plate balance scale (accuracy ± 1 g), was packed into a foamed polystyrene tray. Then, the PP tray filled with pollen paste was wrapped to prevent the evaporation of moisture from the tray and stored in a refrigerator to prevent alteration of the PP. The pesticide-free PP tray was stored in the refrigerator compartment, and the PP tray containing a pesticide was stored in the freezer compartment.

##### 1.2.3 Preparation for field experiment

Field experiments were conducted five times at the resort in mid-west Japan (Shika-machi, Hakui-gun, Ishikawa prefecture, Japan). Four field experiments (T. Yamada et al., 2012; 2018a; 2018b; 2018c) were conducted at the same location, while the remaining one field experiment (T. Yamada, 2020) was conducted at another location nearby. In addition, another field experiment was conducted from 2014 to 2015 in Maui, U.S.A. (T. Yamada & K. Yamada, 2020), and that is, six field experiments were conducted so far. The experimental environment, experimental methods, and publication outlines of four field experiments other than the field experiment conducted in Maui (T. Yamada & K. Yamada, 2020) where there is no cold season and the field experiment conducted in 2010 (T. Yamada et al., 2012) when the high accurate measurement method was not established yet, are summarized (Table S2).

In field experiments to investigate the effects of pesticides on bee colonies, it is necessary to minimize the effects of pesticides other than the pesticides administered in the experiments. Therefore, we tried to make the area around the experimental site as pesticide-free as possible. In other words, in the long-term field experiments in the Midwest of Japan, a water-feeding spot and foraging ground for bees were built in the immediate vicinity of the experimental site. Specifically, a system was installed that automatically supplies fresh water from a large tank. And, in a pesticide-free environment, new plants (rape blossoms *Brassica napus*, hairy vetch *Vicia viliosa Roth.*, white clover *Trifolium repens*, Chinese milk vetch *Astragalus sinicus*, wingstem *Verbesina alternifolia* and so on) and fruit-bearing trees (chestnut *Castanea crenata*, persimmon *Diospyros kaki*, mulberry *Morus alba*, Cherry tree *Prunus avium,* Japanese plum *Prunus salicina*, pawpaw *Asimina trilob*, etc.) were planted and cultivated (T. Yamada et al., 2018a). In addition, a long-term field experiment in the Maui, U.S.A. was conducted in woods of macadamia nuts grown without pesticides (T. Yamada & K. Yamada, 2020).

Here, we will introduce the experimental preparation concretely. Each hive-box was arranged at intervals of 80 cm, with its entrance facing south, on a stand of about 20 cm in height, and then a large PVC tray with many perforations of 3mm in diameter to prevent water from collecting was placed between the hive-box and the stand for the measurement of the number of dead bees.

While keeping in mind that the observation results of the photo images are analyzed, various measures were taken to prevent errors in field experiments. For example, in order to be able to infer the experimental contents of the bee colony in the hive-box, a large display tag was attached at the top of the front of each hive-box. A label was attached to the bottom of the hive-box or on four walls, displaying which part of the hive-box it was. Moreover, several labels displayed so that the position in the hive-box of each comb-frame and the front and back of the comb-frame could be seen were attached to the pier of the comb-frame. In addition, the name of the hive-boxes to be administered was pre-written on the sugar syrup (SS) container and the pollen paste (PP) tray so that the hive-boxes to be administered could be found.

##### 1.2.4 Procedure for field experiment

The most important data in the field experiment of honeybee colony are both the number of adult bees and capped brood, which are indicators of the size, strength, vitality and power of the colony. To obtain these data as accurately as possible, the field experiment was carried out by the following procedure.

**STEP-1)** To avoid being stung by a bee, dress firmly with protective measures, and conduct experiments.

**STEP-2)** Prepare experimental tools, recording paper, containers with SS, trays with PP, camera, the empty hive-box or the like.

**STEP-3)** Take a picture of the experiment site (entire hive-box) before the experiment (check whether abnormal conditions have occurred).

**STEP-4)** Count the number of dead bees in a large tray placed under the hive-box while picking up with tweezers (the number of dead bees outside the hive-box). Throw away what you’ve finished counting out of the tray. It should be noted that the number of dead bees in the hive-box is counted during the internal inspection of the hive-box. The number of dead bees inside and outside the hive-box is summed up and recorded as the number of dead bees on the date of the measurement.

**STEP-5)** Take a picture of the front of the hive-box on whose front the label for displaying the content of the hive-box is attached before the inspection of the inside of hive-box in order to make sure that the experiment target of the photography is not wrong.

**STEP-6)** Open the hive-box and remove the cloth that covered the top of the comb-frame of the hive-box, and take a whole photograph of the top of the hive-box. In addition, gently return the bees attached to the cloth in the hive-box not to prevent them from escaping from the hive-box.

**STEP-7)** The tray with PP and the container with SS (tray, feeding frame), which were put in the hive-box on the previous experimental date, are taken out from the hive-box, measuring the remaining amounts of PP and SS by the upper dish balance. Incidentally, when taking out the containers and the tray from the hive-box, gently return the bees attached to the container in the hive-box.

**STEP-8)** Gently remove the comb-frame with bees from the hive-box in the order of number (in order from left to right toward the front of the hive-box), take a picture of both sides of each comb-frame. When you are taking a picture, if you find a queen bee, after taking a picture of the queen bee, put it in the queen cage and once isolate it. When finishing the photograph of the comb-frame, gently put the comb-frame in the spare empty hive-box prepared beforehand, and the lid of the hive-box is closed. The queen bee in the queen cage is placed at the top of comb-frame in the spare hive-box.

**STEP-9)** After taking pictures of both sides of all comb-frames with bees, take pictures of the four sides and the bottom in the hive-box in order to determine the number of adult bees left in the hive-box. If adult bees are outside the hive-box as well as inside the hive-box (sometimes near the entrance of the hive-box at tropical nights), take a picture to count the number. The sum of these numbers of adult bees are regarded as the number of adult bees of the relevant colony (hive-box).

If you find a queen bee when you’re taking a picture of a comb-frame, take its picture, if you cannot find it, check for the presence or absence of the queen on the comb-frames and inside the hive-box a few times. If the presence of the queen bee cannot be confirmed even after a few-times confirmations, after checking the photographs taken on that experimental date with expanding them, check again the queen bee in that hive-box on the next experiment. If you cannot find a queen bee at the next check, determine the absence or absence of a queen bee in the relevant hive-box after judging from the situations on queen cells, oviposition, the larvae, etc.

**STEP-10** Once the number of adult bees has been measured and the queen has been confirmed, the bees attached to the comb-frame are shaken off from it to make a comb-frame without bees in order to determine the number of capped brood in each bee colony. In this way, the photographs on both sides of the comb frame without bees make it very easy to count the number of capped brood. The specific steps are described below.

**STEP-11)** In Take out the comb-frame with bees that had been placed in the spare hive-box in original numerical order, while gently shaking the bees off the comb-frame in the hive-box, and return all the bees attached to the comb-frame to the original hive-box. After taking pictures of both sides of the comb-frame without bees, return the comb-frame without bees to the original hive-box. Repeat the above process for all comb-frames.

By the way, at the point when you return a few bee-free comb-frames from the spare hive-box to the original hive-box, move the queen-cage containing the queen bee from the top side of the comb-frame of the spare hive-box to the top side of the comb-frame of the original hive-box. Once you have taken photos of both sides of all the comb-frame without bees, and have returned all the comb-frames from the spare hive-box to the original hive-box, remove the queen bee from the queen cage and return it to the original hive-box.

**STEP-12)** When taking a picture of the comb-frame without bee, you may discover some abnormality, such as queen-cells, *Varroa* mites, wax-moth larvae and so on. In this case, keep taking pictures of them. These anomalies are not limited to this situation, and if you find anything, record it in photos as evidence.

**STEP-13)** If there is a suspicion of the absence of the queen bee (death or disappearance, etc.), the division or escape of the bee colony, the attack of the giant hornet, the spread of mites, the infection of chalk disease, foulbrood, etc., the impact on the experiment will be significant, so if you find such an event during experimental observation, record it as a photographic image and record the detailed observation results in the laboratory notebook.

**STEP-14)** After covering the cloth on the top of comb-frames, put new SS and PP in the corresponding hive-box, the measurement is ended after the hive-box is covered by its lid.

**STEP-15)** After completion of the experiment, the test contents are confirmed and described in the laboratory notebook.

#### 1.3 Supplementary Method S3: Counting methods of adult bees, capped brood, mite-damaged bees

Using the photographs taken as described above, it was possible to determine the numbers of adult bees, capped brood and mite-damaged bees. Counting such numbers from photographs is an extremely difficult task. Therefore, with the cooperation of Mr. Yoshiki Nagai of Nanosystem Co., Ltd., Kyoto, Japan (http://nanosystem.jp/firm.htm), which develops image processing software, the author has developed computer software in 2012 that automatically counts the relevant numbers from photographic images, and improvements have been made to the software since. The accuracy and operation of the counting have been greatly improved. By using this software, the time and accuracy for counting such numbers have been significantly improved, compared to the amount of work required to count by hand directly from the original photo.

However, these counts are still inaccurate because of bee overlap, out-of-focus images, extreme contrast differences in the images, misjudgments of the counting targets, and so on. In this software, it is also possible to manually correct after automatic counting. Therefore, after automatic counting, the counting mistakes due to the automation were corrected by manual operation while zooming into the image, and the corrected number became the final data. Thus, errors still emerge in automation when using this software, but by using manual operation, it is possible to correct the counting errors. The improvements in the counting accuracy and speed under this software contributed greatly to the improvement of the data analysis accuracy and the speed in the field experiments. An overview of this automatic counting system is provided below.

##### 1.3.1. Software developed to assist in accurately counting the numbers of adult bees, capped brood, and mite-damaged bees

This software was developed to count the numbers of adult bees and capped brood accurately. The following explains how to count the number of adult bees present in the image of a comb-frame and how to count the number of capped brood using the image read.

The numbers of adult bees and capped brood are counted with software using the image from a photo imported into the computer. The number of adult bees in the colony is obtained by summing the numbers of adult bees on both sides of all the comb-frames with the bees and the number of remaining adult bees in the hive-box without the comb-frame. The number of capped brood in a colony can be obtained by summing the numbers of capped brood on both sides of all the comb-frames without bees. The number of adult bees damaged by the *Varroa* mite can be obtained from the same photo used to count the adult bees. How to count bees damaged by *Varroa* mites is described in a separate section. The measurement results obtained by this method, as long as the image remains, can be verified by anyone at any time. In the following sections, the determination procedure of the numbers of adult bees and capped brood, and that of the number of mite-damaged bees, is described separately.

##### 1.3.2 Determination procedure of the numbers of adult bees and capped brood

**STEP-1)** The photo image is automatically binarized by the software. Next, the area to be measured is set on the image by using the difference in brightness on the image between the place where adult bees exist and that where they do not, the difference in brightness between the place where capped brood exist and that they do not, and the difference in the features of the adult bee and the capped brood. The number of the identified adult bee or the number of the identified capped brood is counted in the area to be measured of the photo image.

**STEP-2)** In order to count as accurately as possible, before the binarization process of the image, divide the entire image into several areas that seem to have the same threshold value in each area (up to 4 areas) as below. Then, enter the optimal threshold for each divided area (the maximum number of divided areas 4, the split shape is optional).

Examples of division: (1) Distinguish between the capped honey area and the capped brood area. (2) Distinguish between areas where there are few bees and areas where bees are dense. (3) Distinguish between different areas of contrast or lightness (e.g., the part where the light hits and the part of the shadow).

**STEP-3)** Enter the type of key to mark counted adult bees and capped brood one by one (+, *, numbers, etc.) and the color and size of key ((you can change them after the end of the count).

**STEP-4)** When the count is performed, while automatically marking those counted bee or capped brood in the image, continue counting, the number of counts at that time is displayed at the bottom of the image. After counting the adult bee or capped brood in the image in a short time, the total count in the image is displayed at the bottom of the image. The count status is visually determined, if dissatisfied, by changing the threshold value, recount. In a few tries, even if you change the threshold, if there is no improvement in the count situation, the count of approximate number by automatic measurement in the computer is assumed to have been completed.

**STEP-5)** After automatic counting by the computer, using the same software, switch to manual count operation, and then correct the count error at the time of automatic counting. Therefore, using a large screen monitor, while further enlarging the image, mark-mistakes of adult bees and capped brood during automatic counting (duplicate count, counting things that are not the object) and forgetting to mark (due to the overlap of bees, the blurred image, the extreme differences in light and dark, etc.) are corrected while visually checking. All possible efforts on this correction are exerted for obtaining the number of adult bees or the number of capped brood as accurate as possible (most important and serious work).

It should be noted that, even when the software is manual counting, by the removal and grant of the mark, automatically change the number of bees or capped brood. You can resume this fix at any time. If it is determined that the correction is complete, it moves to the measurement of the next new image.

**STEP-6)** When posting the measured number of adult bees or capped brood in a separate table, call the measured image, check again in the enlarged image, after checking whether there is a count error (correct if you find a mistake), the number is posted to the separate table.

The sum total of the numbers of adult bees on all combs and those of adult bees on four walls and bottom in the hive-box becomes the number of adult bees in the bee colony at a certain measurement date. In a similar manner, the sum total of the numbers of capped brood of all combs, becomes the number of capped brood in the bee colony.

##### 1.3.3 Counting procedure of the number of mite-damaged bees

Various sampling methods for *Varroa* mites in a honeybee colony, such as the sticky board method, the roll method, and the powder sugar shake method, have been reported (Strange & Sheppard, 2001; Macedo et al., 2002; Barlow & Fell, 2006). With these methods, it is impossible to determine the total number of *Varroa* mites in a bee colony.

It is presumed that the impacts of *Varroa* mites on a bee colony appear directly on the adult bees with damage caused by the mites. This section aims to measure the total number of bees damaged by mites, not the total number of mites in the bee colony.

To count the number of mite-damaged bees as accurately as possible, the images and software used to count the number of adult bees were used again. Using this counting technique for the number of adult bees, it is relatively easy to count *Varroa* mites on an adult bee. On the other hand, it is very difficult to count *Varroa* mites that causes damage to a brood in a comb cell. If any damage is received from *Varroa* mites during the brood stage, there should be a trace of the damage even if the brood becomes an adult bee. An adult bee with traces of its brood stage is considered to be a mite-damaged bee. For example, it is conceivable that a mite may fall out during the development of an adult bee. So, we checked to see if there were any traces of mites left on the adult bees. Therefore, assuming that such traces confirm the presence of mites, we checked the adult bees for traces of mites.

The criteria for the determination of such traces were created based on the information obtained by enlarging the image as is described below. The examples certified to be mite-damaged bees are shown in Figure S5 (T. Yamada, 2020). According to this criterion, using the photographic image taken for the adult number measurement, it is possible to evaluate the situation of the *Varroa*-mite damage among almost the total number of adult bees in the bee colony.

Therefore, the total number of both adult bees with *Varroa* mites and those with traces of *Varroa*-mite damage was determined manually with the software used to measure the number of adult bees while enlarging the adult bee number measurement image. The measurement of the number of mite-damaged bees was a very difficult task and took considerable time (about 6 months). To eliminate the errors between operators, this measurement was carried out by only one person. Figure S6 shows the seasonal changes of mite-damaged bees in a bee colony measured according to the following criteria.

##### 1.3.4 Criteria for determining whether or not it is a mite-damaged bee

Just before capping a comb cell where a larva lives, female *Varroa* mites enter into the comb cell, and then lay eggs and multiply in it. Therefore, we determined that it would be impossible to count the number of mites infesting bee-colony, directly.

In this study, rather than a method of directly counting the number of *Varroa* mites in the bee colony, it is assumed that the number of bees damaged by mites is equivalent to the degree of the bee colony damage caused by mites. The number of mite-damaged bees on the enlarged photographic image of the comb with bees taken to measure the number of adult bees, using the software developed to count the numbers of adult bees and capped brood mentioned above, is manually counted while referring to Figure S5 according to the following criteria.

In this method, it is not possible to directly evaluate the absolute number of mites in the bee colony, but it is possible to estimate indirectly the damage to the bee colony caused by mites. Additionally, since a photographic image for the measurement of the number of adult bees in a field experiment is used to evaluate the damage to the bee colony caused by *Varroa* mites, it is possible to re-measure the number of mite-damaged bees while checking again and again whenever you need, and it is possible to improve the measurement accuracy. Further, using the photographic image for the past adult number measurement, it is also possible to measure the number of mite-damaged bees during past experiments.

Notes and criteria for determining whether the damaged bee by the *Varroa* mite using the captured image is shown below.

Female *Varroa* mites enter the comb-cell where the larva is just before capping the cell, lay the egg (all male mites) and the number of mites increases in the capped comb-cell. Not only the larvae, it is said to continue to cause damage to adult bees. However, it is impossible to count the number of mites in the capped comb-cell. Therefore, although indirect, adult bees with the traces of mite-damages (the presence of mites can be confirmed, there are traces of swelling and inflammation that seem for the bee to have been bitten by mites, wings of the bee are deformed or fallen out, there is a missing mark of the mite on the bee) is here regarded as mites. That is, the number of adult bees having the above traces can be regarded as an index to estimate the degree of the mite-damages of the bee colony.

By the way, there are cases when adult bees have not only traces of mites, but also mites themselves. Of course, in that case, count the number of mites attached to the adult bees.

The specific criteria for judgment are shown below. It is suspected that the three-dimensional one attached to a bee, which is small and round (including the ellipse), is a mite in the color of the red-brown system (including the oval) and red-brown system (including the semi-transparent). A suspected mite is determined whether it is a mite based on the criteria below described and Figure S5, while being examined from various angles, using an enlarged image. If a suspected mite cannot be determined by all means, it is considered a mite or a trace of the mite.

1) What appears to be planar rather than three-dimensional in the vicinity of the neck is considered to be an innate pattern of a bee appearing when the bee bends its neck or a shadow of other bee wings, etc., and is not considered damage by mites.

2) Tiny stuff with colors attached to adult bees other than red-brown and black (especially whitish ones) are regarded as pollen.

3) What the wings appear three-dimensional like wrinkles deformed is regarded as a shadow due to the shrinkage of the wings. However, the cause of the shrinkage of the wings is a large possibility of deformed wing virus mediated by *Varroa* mites. The state of the shrinkage of the wing is scrutinized on the image to determine whether the shrinkage is due to mites.

4) If a bee with shrunk or missing wings has traces of a mite bite, it is considered a mite-damaged bee, even if it is not possible to confirm the presence of a mite.

5) If you can’t distinguish between mites and pollen or garbage, you should consider them mites. However, if it is clear that it is other than mites such as pollen and garbage, exclude it.

6) Planar ones are usually determined to be not mites. However, after scrutiny, exceptionally, it may be considered a mite-damaged bee.

7) Except for those that seem to be the rise of muscles, etc.

8) The difference between a mite and garbage is judged by shape, color, surrounding situation, etc., and it counts unnatural things like mites (Subjective, but count even what looks like a mite).

9) Traces bitten by mites (swelling by the bites) are also thought to be damage to bees.

10) What appears to be a mite, regardless of the number of traces bitten by mites, is treated as one mite-damaged bee.

11) Traces of a mite fallen out from a bee are treated as a mite-damaged bee.

Just before the larva’s cell is capped, the mites lay eggs in the food in the cell, since the mites hatched from the eggs grow while sucking the body fluid of the bee’s larva in the capped cell, it is almost impossible to find almost a mite even in the open cell. Therefore, it was decided not to evaluate the mites in the cells.

Mites are present in various places in the hive-box, shape is also easy to mistake for garbage and pollen, etc., also present in the invisible place (especially in the comb-cell), although the absolute number of mites is not known, this method to evaluate by the number of adult bees damaged by mites is believed to be a qualitative indicator of the number of presence of mites. Although the number of mites in the cell cannot be known, the infestation state of mites in the bee colony will be reflected in the number of mites adhering to the adult bee.

This method includes an uncertain element, under the above criteria, by measuring in the same person, and relative comparison of the number of mites between each bee colony, to some extent the time course of each bee group it is considered to represent. By the way, whenever a doubt occurs, again, using the image for the adult bee number measurement taken, recount.

### 2 Supplementary Figures

**Supplementary Figure S1.**
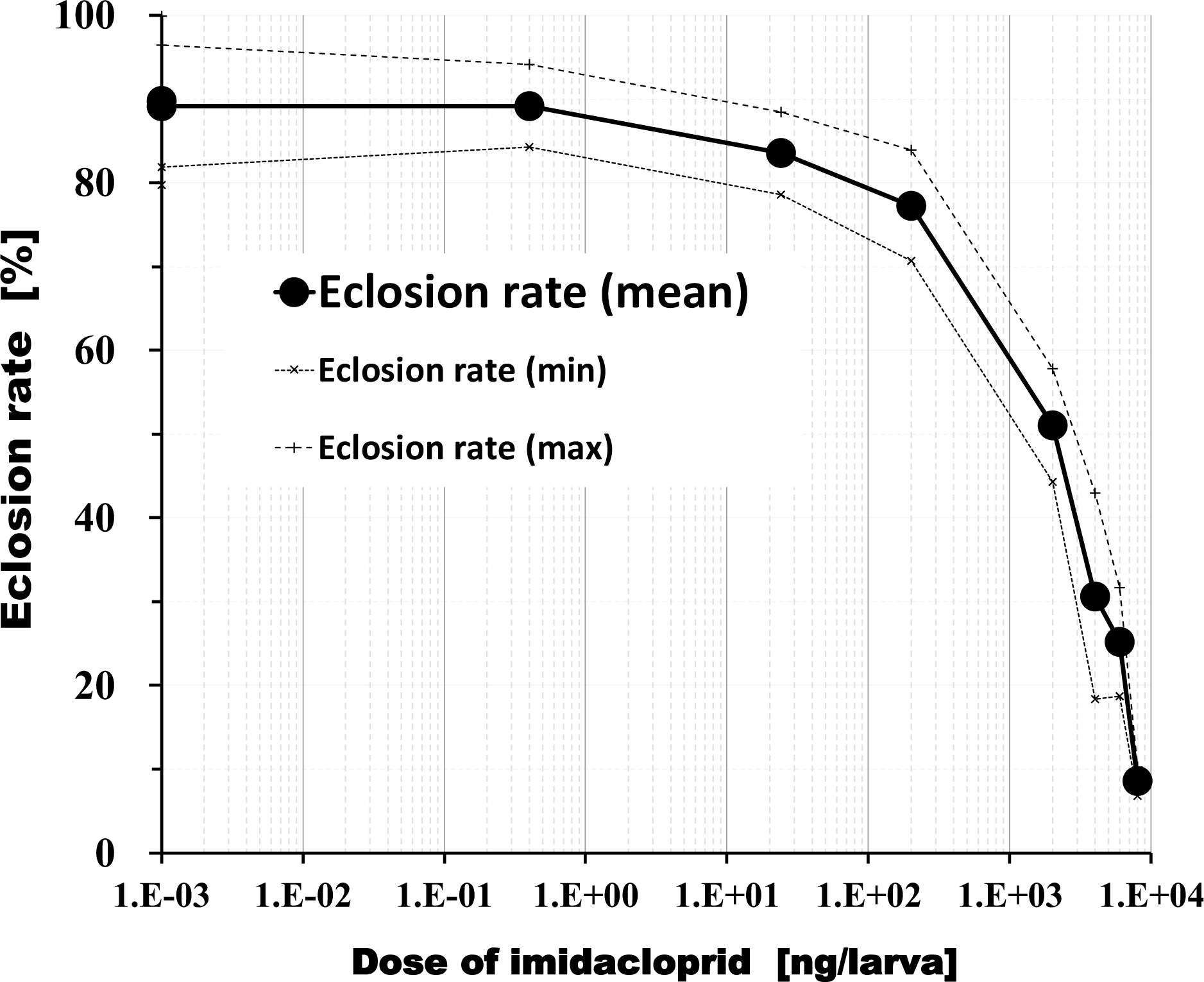
Change in eclosion rate of honeybees with imidacloprid dosage. The source data on the relation between dose of imidacloprid and eclosion rate of honeybee are as follows (Yang et al. 2012): Eclosion rates are; 89.82±10.10 and 89.17±7.26 % for the control group; the rates are 89.20±4.90 % at the imidacloprid concentration of 0.4 ng/larva, 3.54±4.92 at that of 24 ng/larva, 77.29±6.64% at that of 200 ng/larva, 51.04±6.78 at that of 2000 ng/larva, 30.63±12.31 % at that 4000 ng/larva, 25.21±6.47 % at that of 6000 ng/larva and 8.54±1.72 % at that of 8000 ng/larva, respectively. Since Semi-log graphs cannot handle zero values, the pesticide concentration of the control group was set to 0.001 ng / larva for convenience.

**Supplementary Figure S2.**
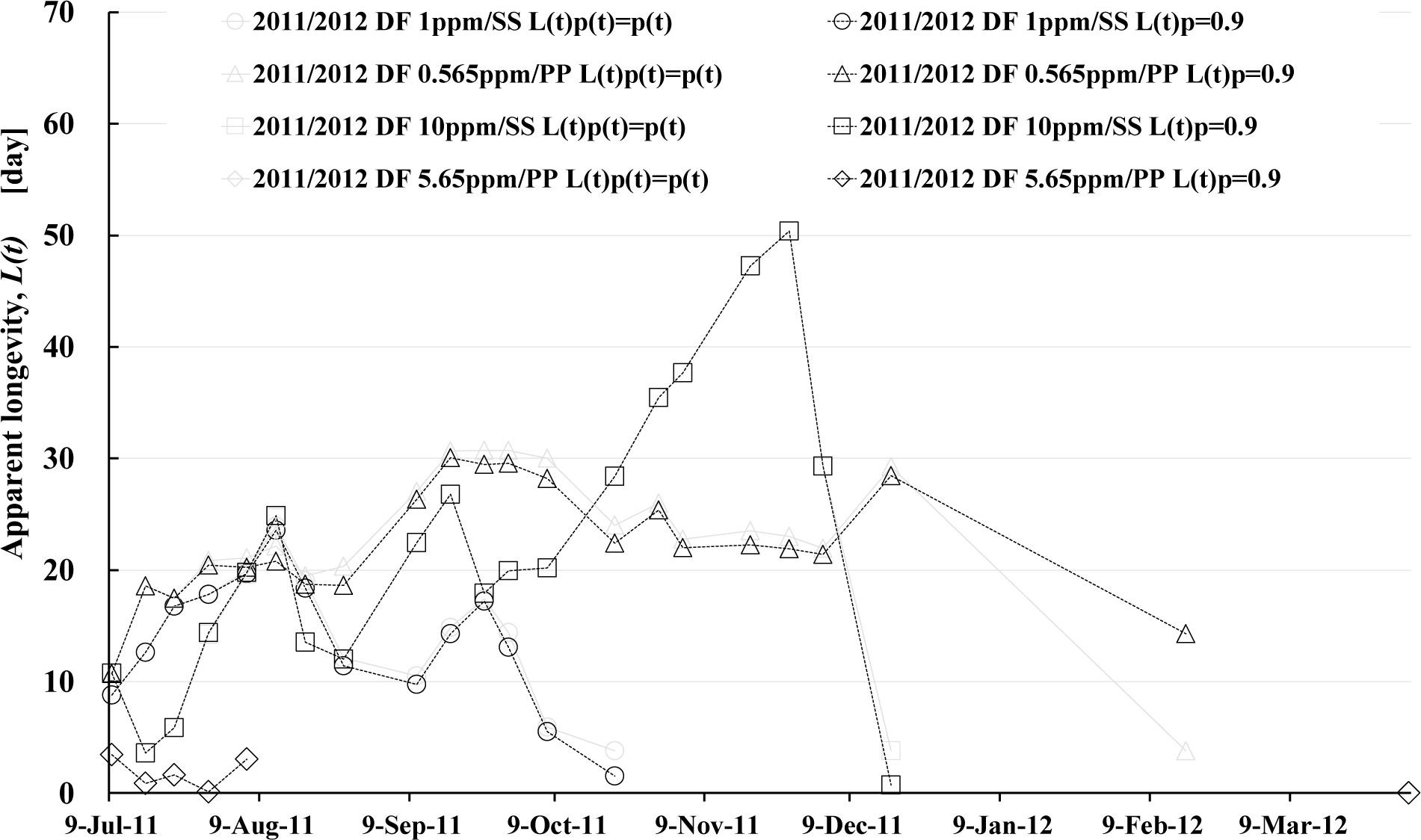
Comparison of apparent longevity between a variable eclosion rate, *p(t)*=*f*(dinotefuran-intake per bee), and a constant one, *p(t)*=0.9. To estimate the apparent longevity of honeybee colony were used the numbers of adult bees and capped brood measured in the field experiment conducted in 2011 to 2012 (T. Yamada et al., 2018a). *L(t)_p(t)=p(t)_* shows the apparent longevity was calculated with the variable *p(t)* changing with the pesticide intake per bee in a period between two consecutive measurement dates; where the ranges of *p(t)* are 0.79–0.9 for the “2011/2012 DF 1ppm/SS *L(t)_p(t)=p(t)_*” colony; 0.84–0.9 for the “2011/2012 DF 0.565ppm/PP *L(t)_p(t)=p(t)_*” colony, 0.8–0.9 for the “2011/2012 DF 10ppm/SS *L(t)_p(t)=p(t)_*” colony, 0.835–0.9 for the “2011/2012 DF 5.65ppm/PP *L(t)_p(t)=p(t)_*” colony, respectively. Incidentally, the intake of dinotefuran (DF) by one honeybee in a period between the observation interval was obtained according to the procedures reported in the previous paper (T. Yamada et al., 2018a). *L(t)_p(t)=0.9_* shows the apparent longevity was calculated with the constant *p(t)* of 0.9.

**Supplementary Figure S3.**
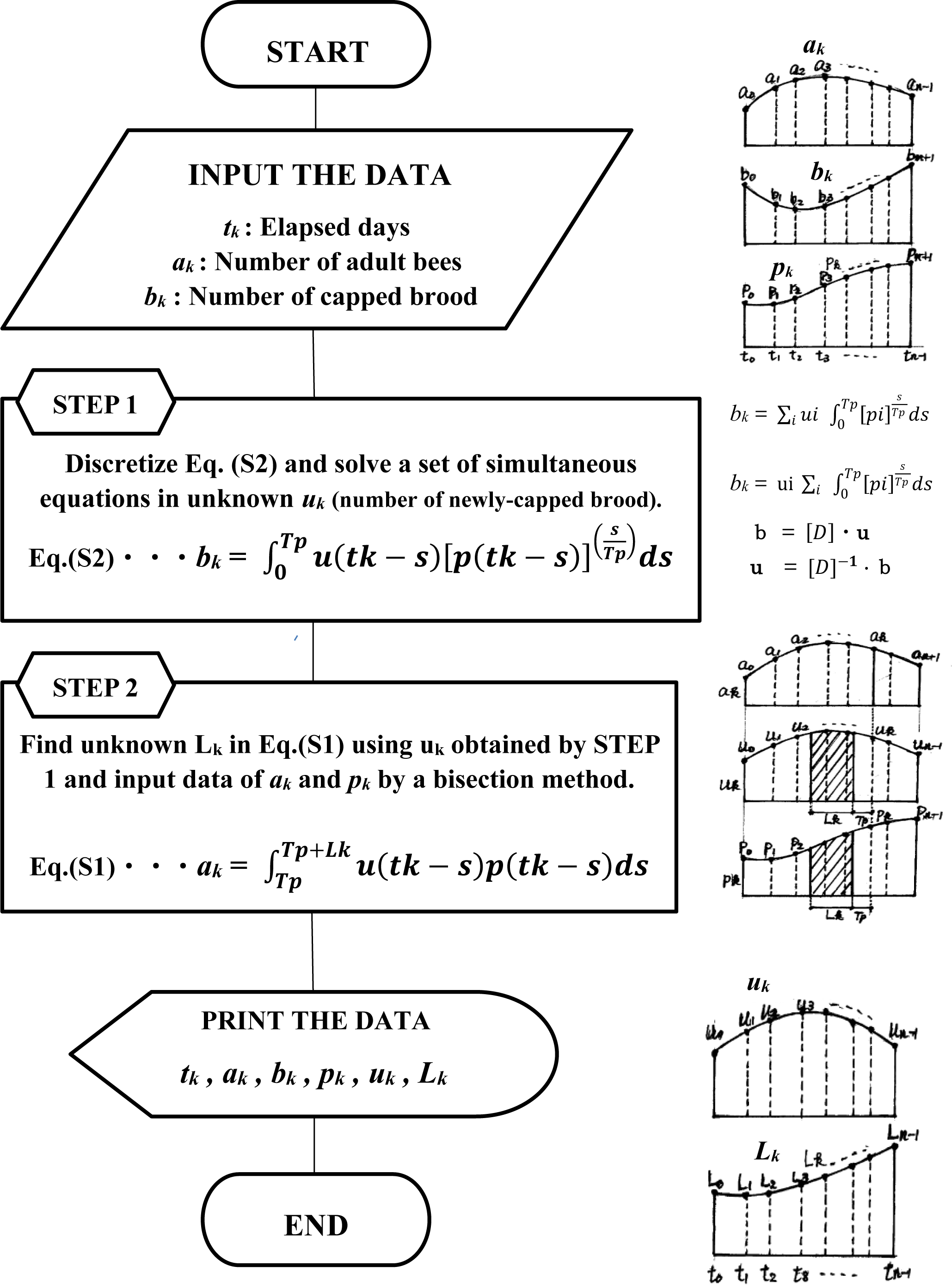
Flowchart to find the apparent longevity, *L(t)*: Mainstream scheme. The calculations were carried out using a software written in the Ruby programming language.

**Supplementary Figure S4.**
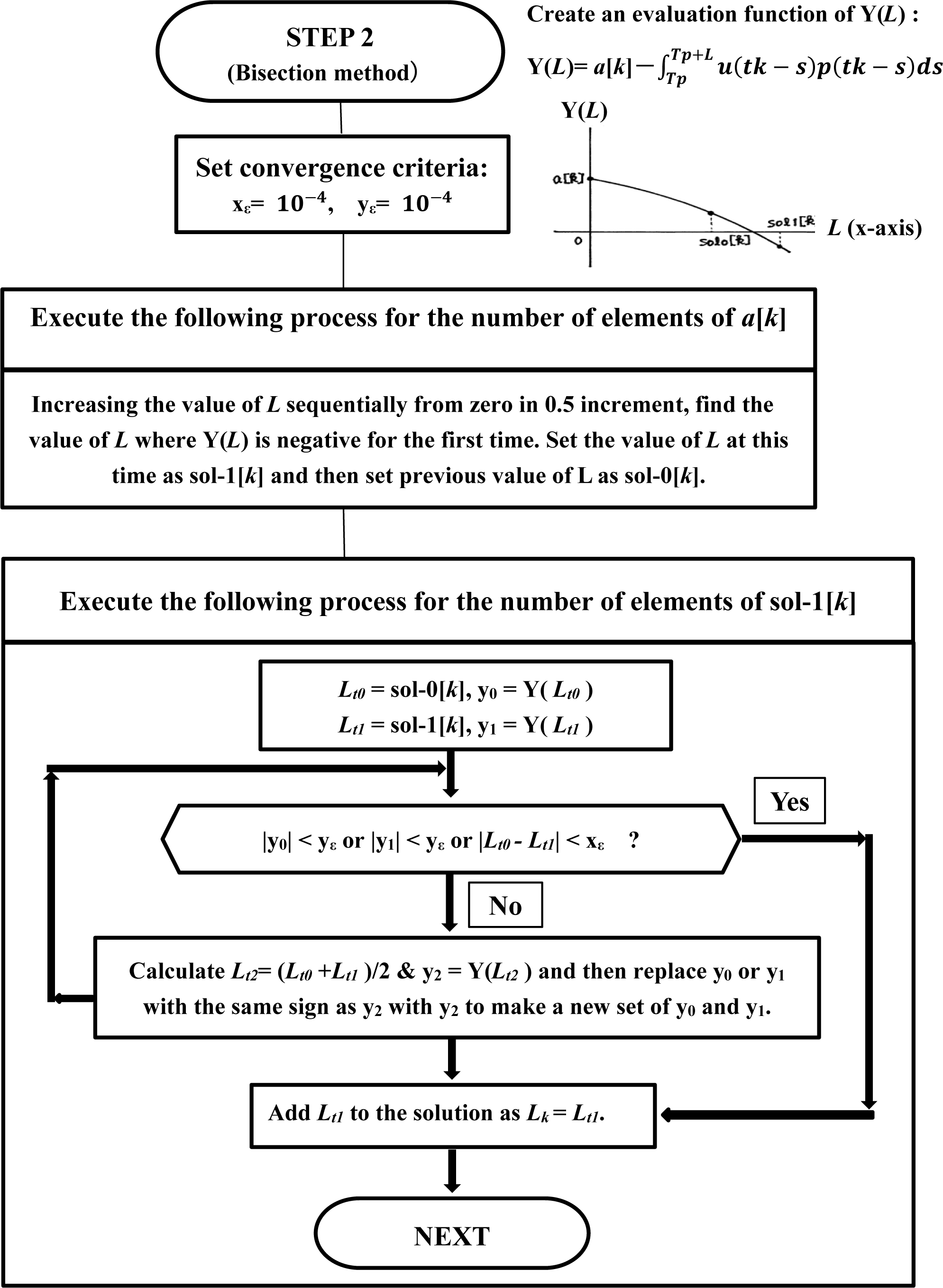
Flowchart to find the apparent longevity, *L(t)*: Bisection method.

**Supplementary Figure S5.**
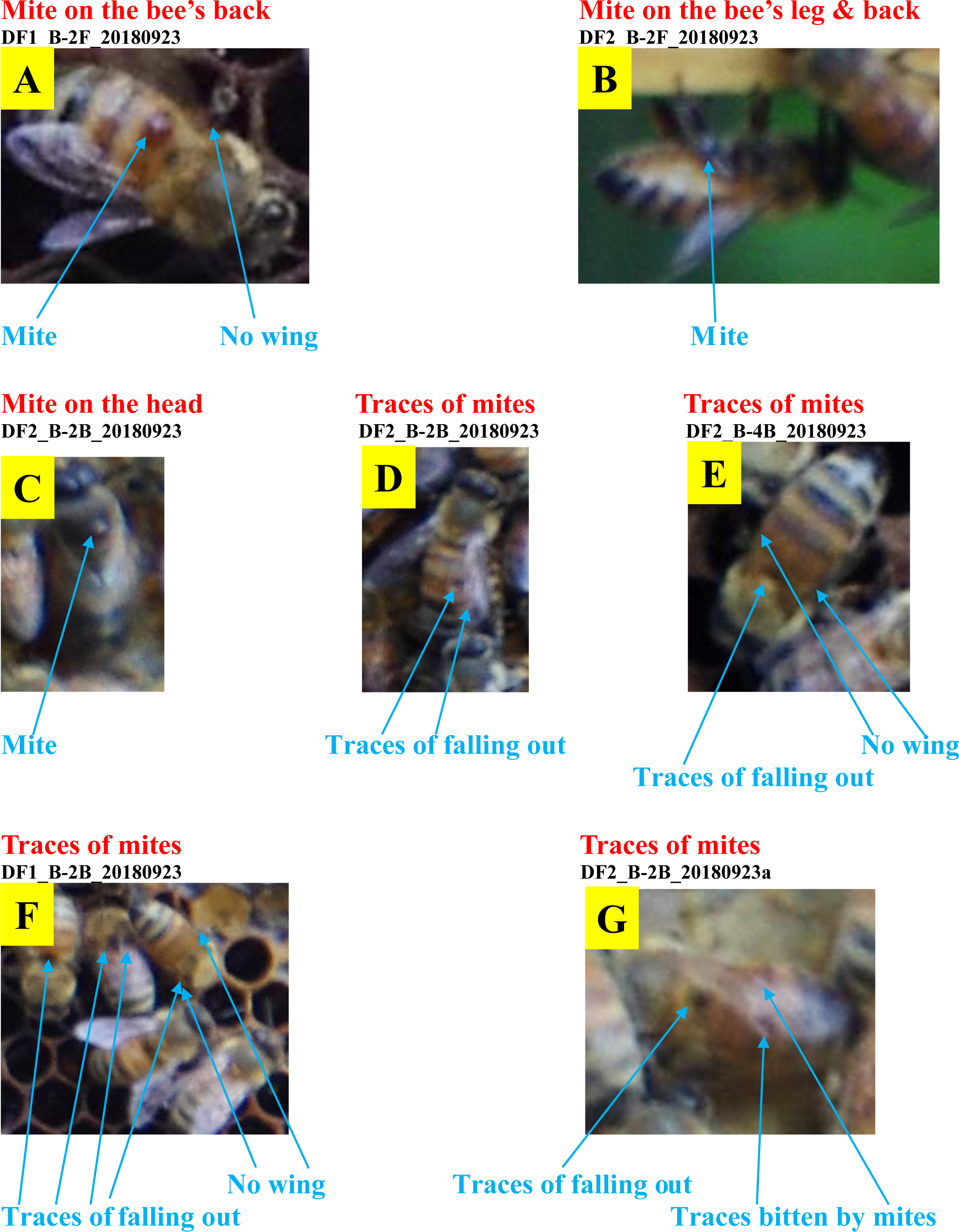
Images of mite-damaged bees. The above images were obtained from the photographs of adult bees on comb-frame taken in the field experiments. Referring the above, we judged whether the adult bee was damaged by mites while enlarging the image. Several representative images of adult bees that appear to have been damaged by *Varroa* mites are shown. Each image is an example of a mite-damaged bee. **A:** A mite on the back of an adult bee (worker bee) whose wing is removed. **B:** Mites on the bee’s leg and back. **C:** A mite on the head of a drone. **D:** The trace of a mite that fell out of the bee’s back. **E:** Trace of a mite that fell out and a bee that has no wings. **F:** Traces of mites that fell out of the bee’s back and a bee with no wing. **G:** Traces of a mite that fell out and a bee that seems to have been bitten by mites.

**Supplementary Figure S6.**
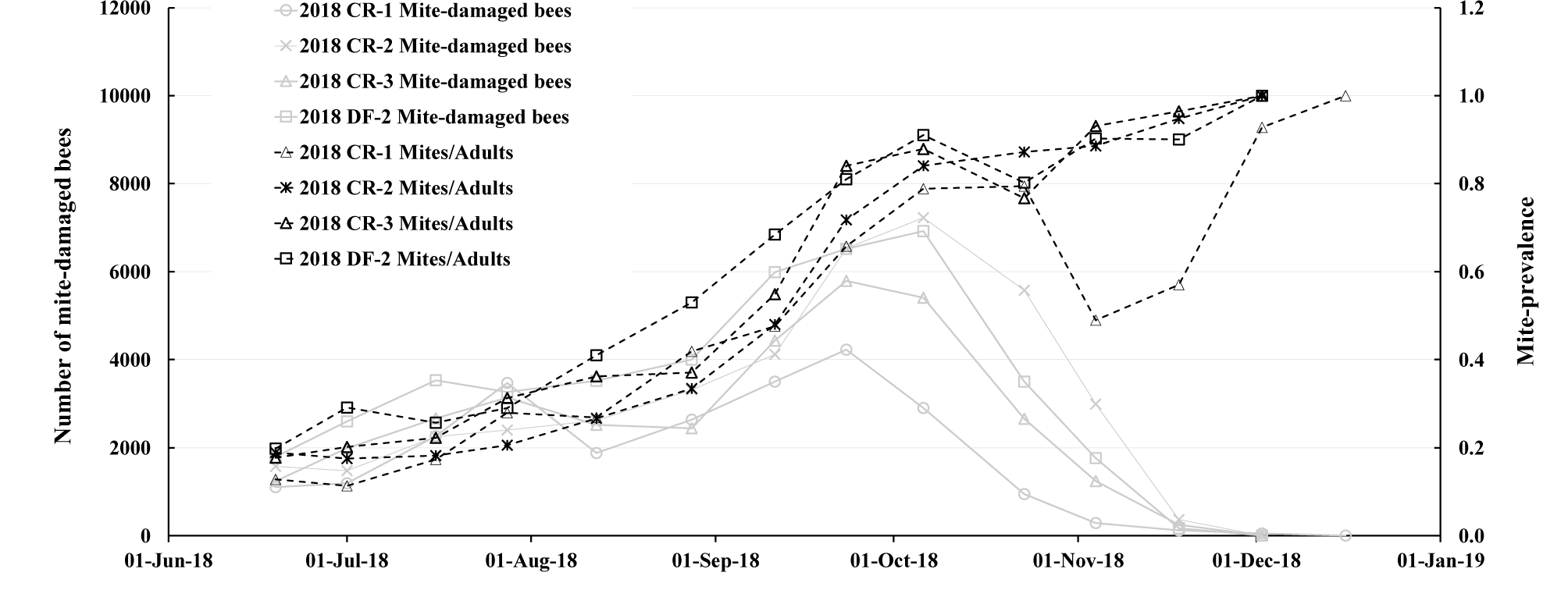
Number of mite-damaged bees and mite-prevalence. Details of these data in the field experiment conducted in 2018 are reported in the previous report (T. Yamada, 2020). The mite-prevalence denotes the ratio of the number of damaged bees by *Varroa* mites (mite-damaged bees) to the total number of adult bees in a period between each measurement interval. Mites/Adults denotes the mite-damaged bees /total bees (mite-prevalence). 2018 CR-1, 2018 CR-2 and 2018 CR-3 denote the control colony which is infested with *Varroa* mites. 2018 DF-1, 2018 DF-2 and 2018 DF-3 denote the colony being infested by *Varroa* mites and to which dinotefuran is administered via pollen paste.

**Supplementary Figure S7.**
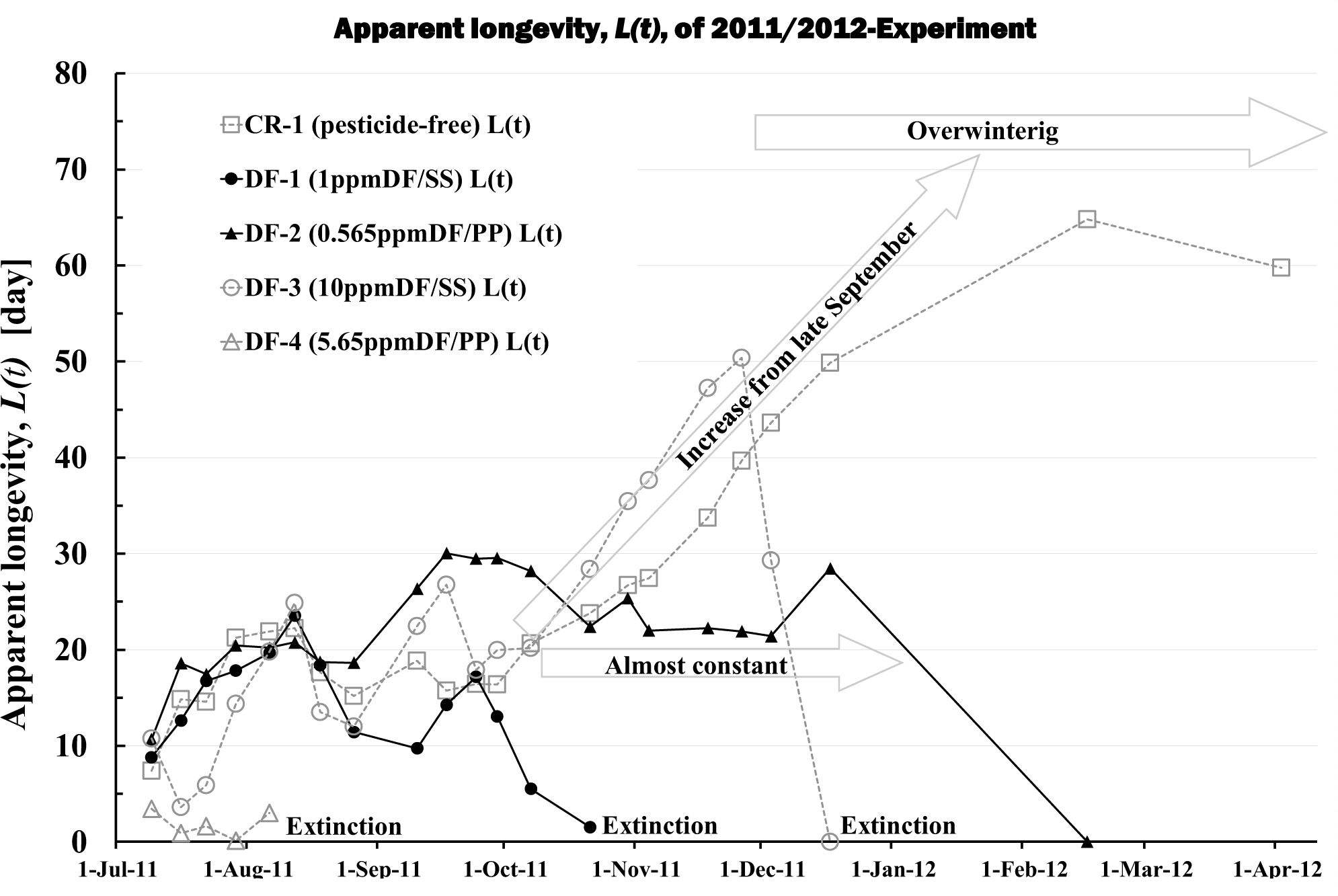
Apparent longevity of 2011/2012-Experiment. **CR-1 (pesticide-free) L(t):** Apparent longevity of control colony (**CR-1**) to which pesticide is not administered. **DF-1 (1ppmDF/SS) L(t):** Apparent longevity of experimental colony (**DF-1**) to which dinotefuran (a kind of neonicotinoids) of 1 ppm is administered via sugar syrup. **DF-2 (0.565ppmDF/PP) L(t):** Apparent longevity of experimental colony (**DF-2**) to which dinotefuran 0.565 ppm is administered via pollen paste. **DF-3 (10ppmDF/SS) L(t):** Apparent longevity of experimental colony (**DF-3**) to which dinotefuran of 10 ppm is administered via sugar syrup. **DF-4 (5.65ppmDF/PP) L(t):** Apparent longevity of experimental colony (**DF-4**) to which dinotefuran 5.65 ppm is administered via pollen paste. All colonies are almost mite-free. The control colony, **CR-1**, and experimental colony, **DF-3**, begin to increase the apparent longevity from late September, as if they already have known that winter is approaching. **CR-1** continues to increase during wintering, but **DF-3** have become extinct soon after wintering. The extinction of **DF-3** is thought to be due to the fact that the colony has been already weakened by dinotefuran before wintering, as evidenced by the fact that other experimental colonies, **DF-1** and **DF-4**, became extinct before wintering. **DF-1** and **DF-3** are both bee colonies that received dinotefuran via sugar syrup, but **DF-1** with a low concentration of dinotefuran became extinct earlier than **DF-3** with a high concentration of dinotefuran (see the previous paper (T. Yamada et al., 2018a) for details). On the other hand, as if another experimental colony, **DF-2**, does not notice that winter is approaching, its apparent longevity does not begin to increase from late September, and the longevity remains almost constant even into wintering. The almost constant longevity changes through the year for a bee colony are quite different from general seasonal longevity changes. This unusual change cannot be explained by the normal physiology of bees. The difference between the seasonal changes in the apparent longevity of DF-2 and that of DF-3 as described above is thought to be due to the difference in the food (vehicles) via which dinotefuran is administer. The mechanism by which this difference occurs is discussed in this paper.

**Supplementary Figure S8.**
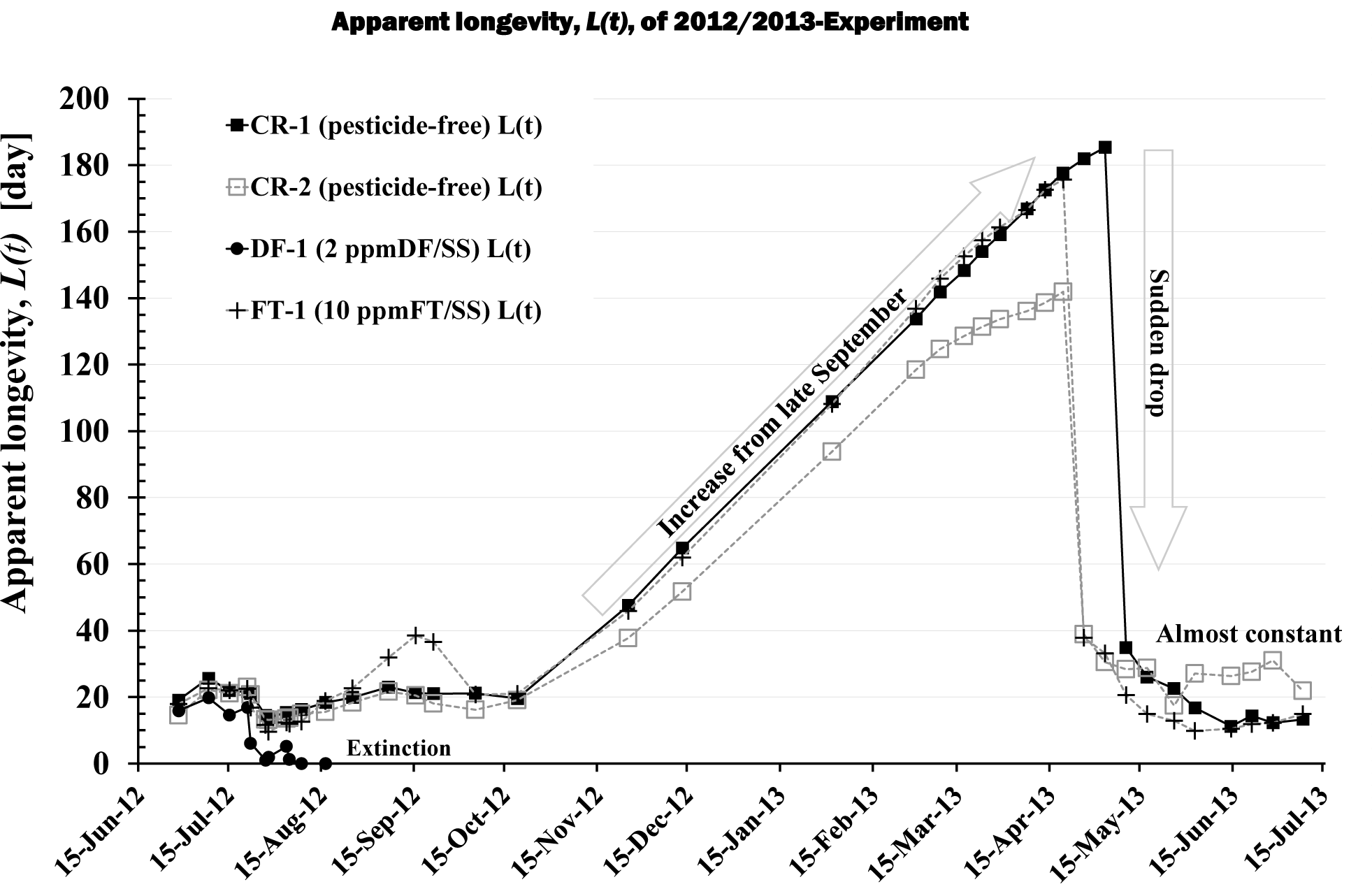
Apparent longevity of 2012/2013-Experiment. **CR-1 (pesticide-free) L(t) and CR-2 (pesticide-free) L(t):** Apparent longevity of control colonies (**CR-1**, **CR-2**) to which pesticide is not administered. **DF-1 (2ppmDF/SS) L(t):** Apparent longevity of experimental colony (**DF-1**) to which dinotefuran of 2 ppm is administered via sugar syrup. **FT-1 (10ppmFT/SS) L(t):** Apparent longevity of experimental colony (**FT-1**) to which fenitrothion of 10 ppm is administered via sugar syrup. All colonies are almost mite-free. The experimental colony **DF-1** became extinct early. Pesticide-free control colonies **CR-1** and **CR-2**, as well as fenitrothion (organophosphate) containing colonies **FT-1**, exhibit general seasonal longevity changes. That is, their longevities are short and almost constant from the end of wintering to late September, then begins to extend from late September, continues to extend until just before the end of wintering, and when wintering is over, it suddenly drops to a short level, after which it stabilizes. It is noteworthy that **FT-1** where is administered fenitrothion of an amount having the same acute insecticidal activities as 2 ppm dinotefuran, succeeded in wintering, just like the control groups (**CR-1** and **CR-2**). The difference between the early extinction of **DF-1** and the successful wintering of **FT-1** is probably due to the difference in the length of the residual effect period. For details, please refer to the previous report (T. Yamada et al., 2018b).

**Supplementary Figure S9.**
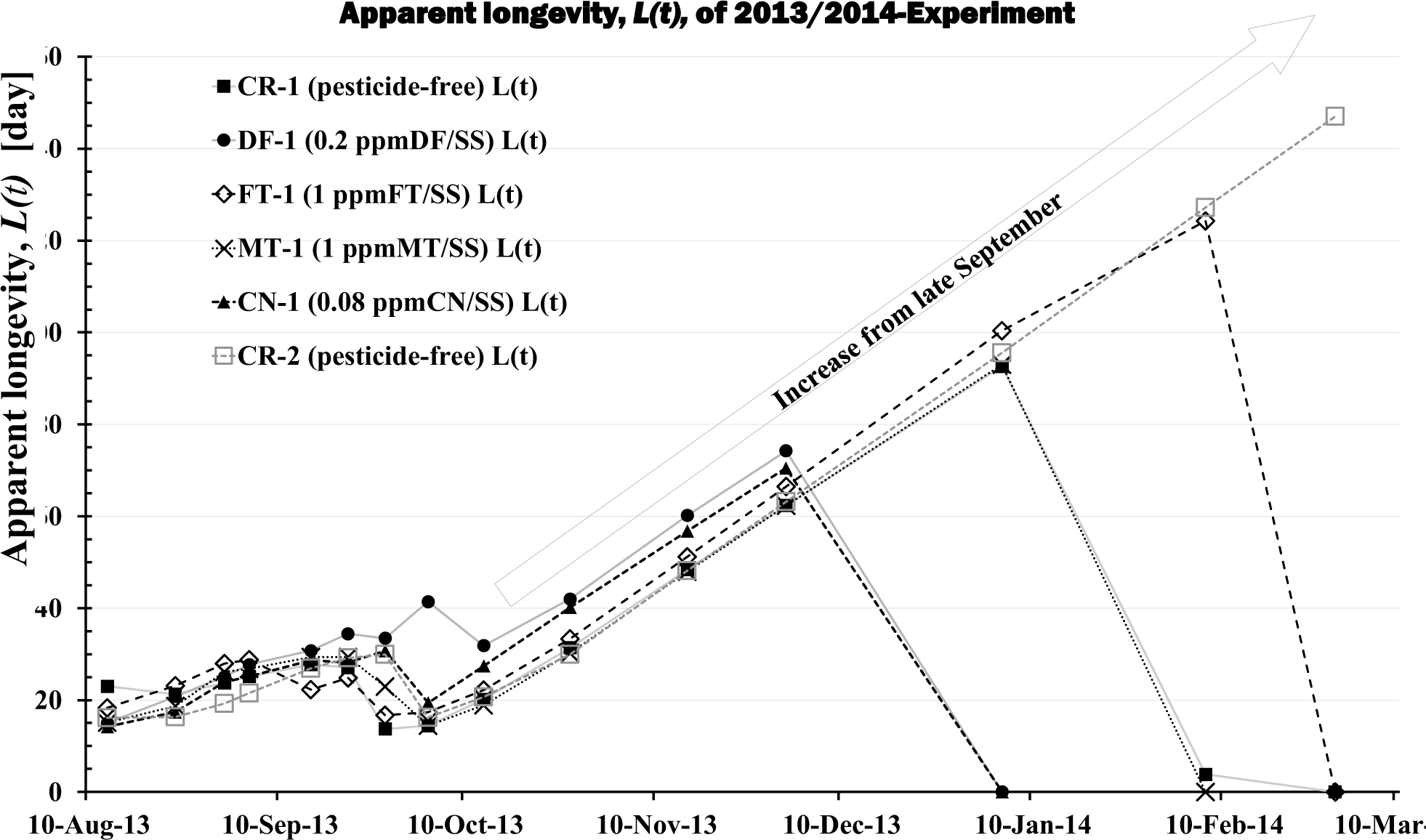
Apparent longevity of 2013/2014-Experiment. **CR-1 (pesticide-free) L(t) and CR-2 (pesticide-free) L(t):** Apparent longevity of control colonies (**CR-1**, **CR-2**) to which pesticide is not administered. **DF-1 (0.2 ppmDF/SS) L(t):** Apparent longevity of experimental colony (**DF-1**) to which dinotefuran of 0.2 ppm is administered via sugar syrup. **FT-1 (1 ppmFT/SS) L(t):** Apparent longevity of experimental colony (**FT-1**) to which fenitrothion of 1 ppm is administered via sugar syrup. **MT-1 (1 ppmMT/SS) L(t):** Apparent longevity of experimental colony (**MT-1**) to which malathion of 1 ppm is administered via sugar syrup. **CN-1 (1 ppmCN/SS) L(t):** Apparent longevity of experimental colony (**CN-1**) to which clothianidin of 0.08 ppm is administered via sugar syrup. All colonies are almost mite-free. The apparent longevity of all colonies (**CR-1**, **CR-2**, **DF-1**, **FT-1**, **MT-1**, **CN-1**) changed in the same way; it held almost constant at a short level from the start of experiment till late September, then began to extend toward winter, and continued to extend during wintering, but all colonies other than **CR-2** became extinct during wintering.

**Supplementary Figure S10.**
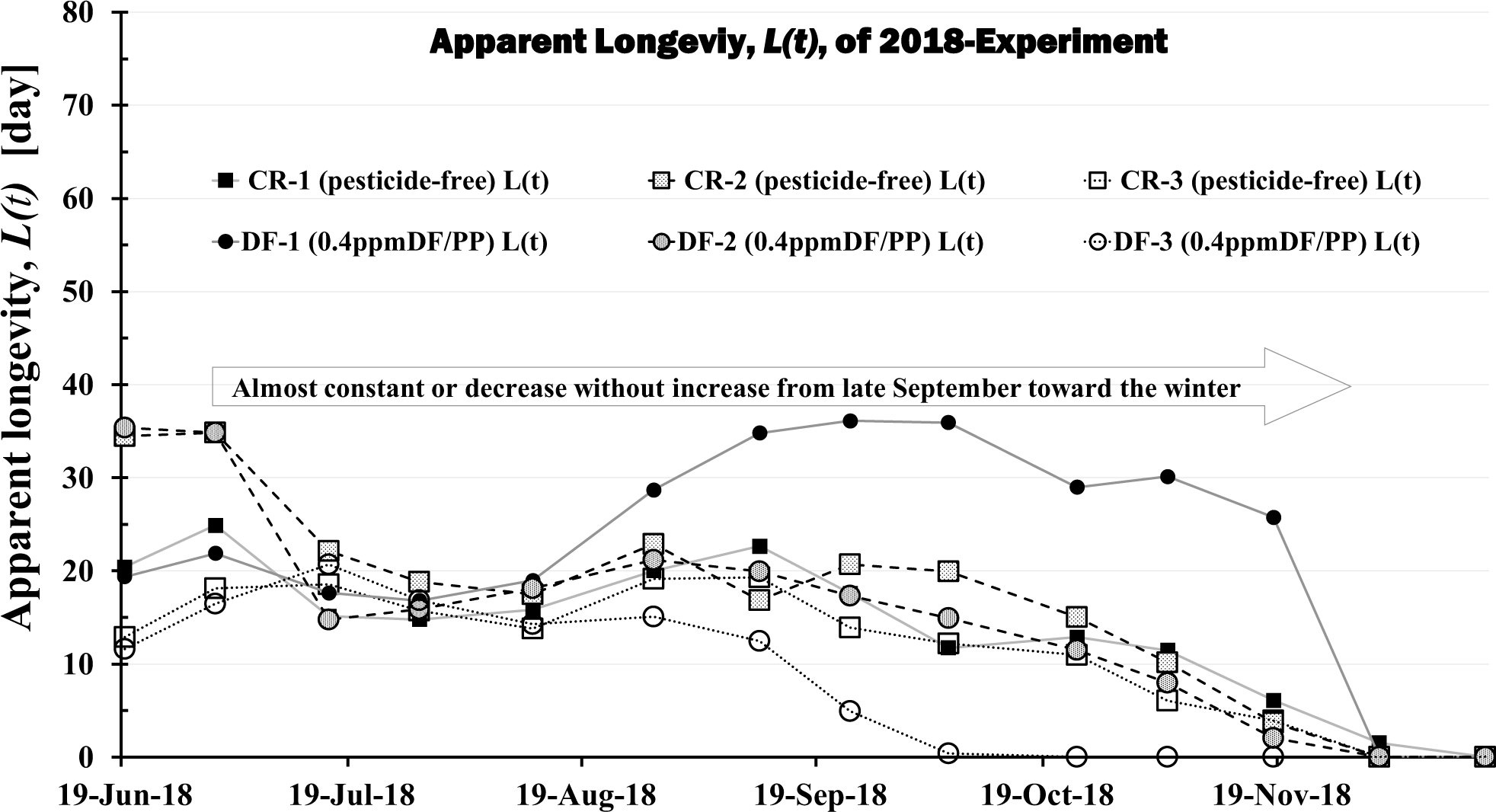
Apparent longevity of 2018-Experiment. **CR-1 (pesticide-free) L(t), CR-2 (pesticide-free) L(t) and CR-3 (pesticide-free) L(t):** Apparent longevity of control colonies (**CR-1**, **CR-2**, **CR-3**) to which pesticide is not administered. **DF-1 (0.4 ppmDF/PP) L(t), DF-2 (0.4 ppmDF/PP) L(t) and DF-3 (0.4 ppmDF/PP) L(t):** Apparent longevity of experimental colonies (**DF-1**, **DF-2**, **DF-3**) to which dinotefuran of 0.4 ppm is administered via pollen paste. All colonies were infested with mites (T. Yamada, 2020). The apparent longevities of all colonies in this field experiment are not the typical seasonal changes that begin to extend from late September and continues until the end of wintering, but has an unusual seasonal change that is almost constant or slightly shorter towards winter. Interestingly, the apparent longevity of all colonies in this experiment is, roughly speaking, fairly constant throughout the year, with little seasonal changes in apparent longevity. This is similar to the seasonal changes in the apparent longevity of bee colonies fed dinotefuran (neonicotinoid pesticides) via pollen paste (T. Yamada et al., 2018a).

### 3 Supplementary Tables

**Supplementary Table S1.** Outline of pesticide intake in three field experiments. Location of experimental site: mid-west Japan (Shika-machi) (Latitude 37°1’9”N; Longitude 136°46’14”N).

|  | Initial number of adult bees | Initial number of capped brood | Administration period of pesticide | Total intake of pesticide per colony during administration [mg/colony] | Total number of adult bees during administration | Total intake of pesticide per bee during administration [ng/bee] | Extinction or survival |
| --- | --- | --- | --- | --- | --- | --- | --- |
| 2011/2012 CR-1 (pesticide-free)<sup>25)</sup> |  |  |  | 0 |  | 0 | Survival |
| 2011/2012 DF 1ppm/syrup<sup>25)</sup> | 2965 | 4263 | 9-Jul-11 to 21-Oct-11 | 4.208 | 13543 | 310.7 | Extinction |
| 2011/2012 DF 0.565ppm/pollen<sup>25)</sup> | 2158 | 2556 | 9-Jul-11 to 3-Dec-11 | 1.8692 | 30779 | 60.7 | Extinction |
| 2011/2012 DF 10ppm/syrup<sup>25)</sup> | 3296 | 3880 | 9-Jul-11 to 16-Jul-11 | 1.9 | 6344 | 290.3 | Extinction |
| 2011/2012 DF 5.65 ppm/pollen<sup>25)</sup> | 1659 | 6993 | 9-Jul-11 to 16-Jul-11 | 0.5257 | 3078 | 65.1 | Extinction |
| 2012/2013 CR-1 (pesticide-free)<sup>26)</sup> |  |  |  | 0 |  | 0 | Survival |
| 2012/2013 CR-2 (pesticide-free)<sup>26)</sup> |  |  |  | 0 |  | 0 | Survival |
| 2012/2013 DF 2ppm/syrup<sup>26)</sup> | 9173 | 9442 | 21-Jul-12 to 16-Aug-12 | 1.55 | 16524 | 93.8 | Extinction |
| 2012/2013 FT 10ppm/syrup<sup>26)</sup> | 8943 | 8732 | 21-Jul-12 to 16-Aug-12 | 17.97 | 19792 | 862.5 | Survival |
| 2013/2014 CR-1 (pesticide-free)<sup>27)</sup> |  |  | 0 | 0 |  | 0 | Extinction |
| 2013/2014 CR-2 (pesticide-free)<sup>27)</sup> |  |  | 0 | 0 |  | 0 | Survival |
| 2013/2014 DF 0.2ppm/syrup<sup>27)</sup> | 6017 | 2214 | 5-Sep-13 to 1-Dec-13 | 0.514 | 8612 | 56.7 | Extinction |
| 2013/2014 CN 0.08ppm/syrup<sup>27)</sup> | 7293 | 1506 | 5-Sep-13 to 1-Dec-13 | 0.4016 | 10130 | 39.6 | Extinction |
| 2013/2014 FT 1ppm/syrup<sup>27)</sup> | 10296 | 3232 | 5-Sep-13 to 28-Feb-14 | 3.86 | 19585 | 197.1 | Extinction |
| 2013/2014 MT 1ppm/syrup<sup>27)</sup> | 7676 | 1396 | 5-Sep-13 to 7-Feb-14 | 6.75 | 15724 | 429.3 | Extinction |

**Supplementary Table S2.**
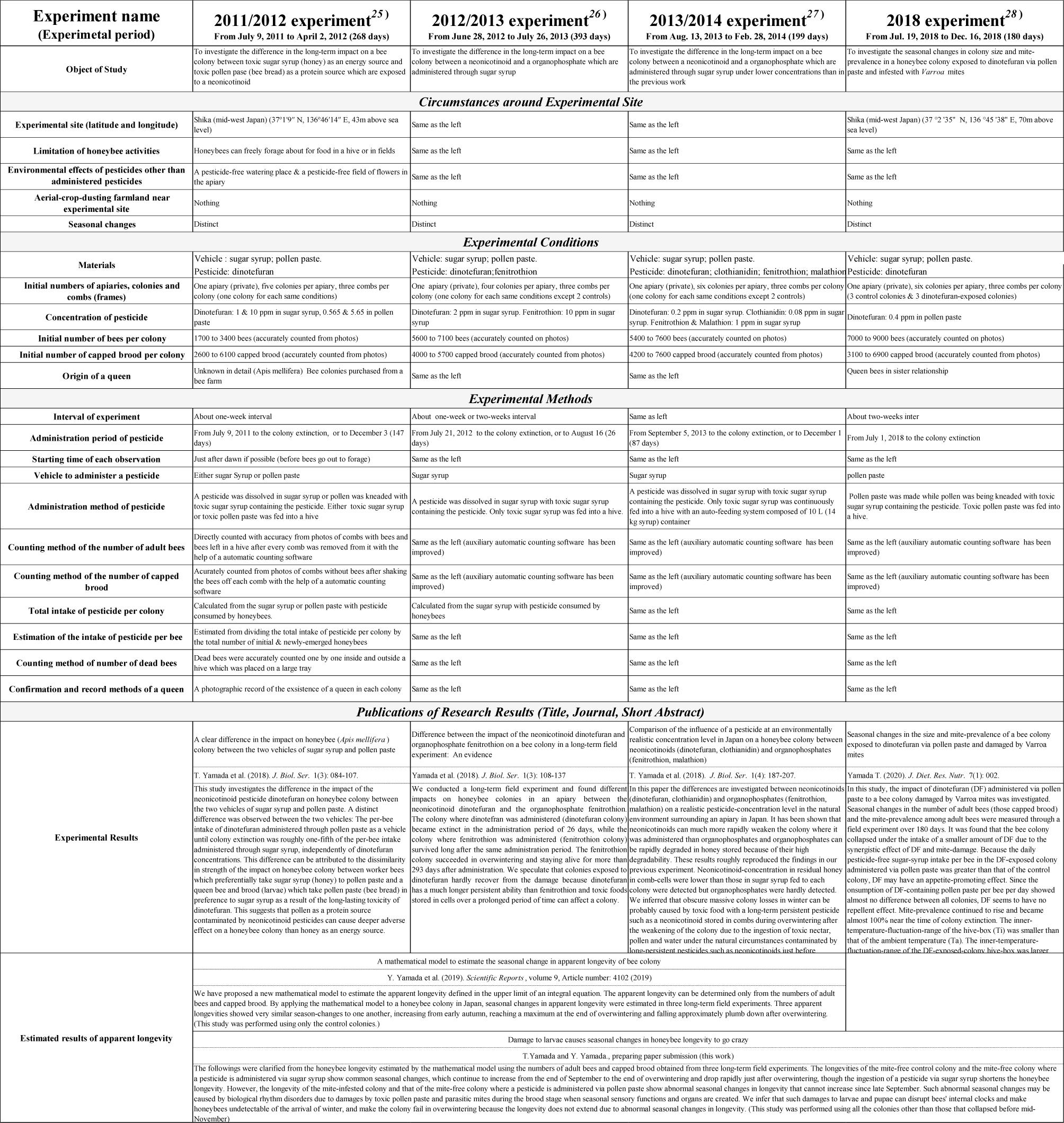
The essentials of experimental conditions and results for four long-term field experiments conducted in Shika, Japan.

**Supplementary Table S3.**
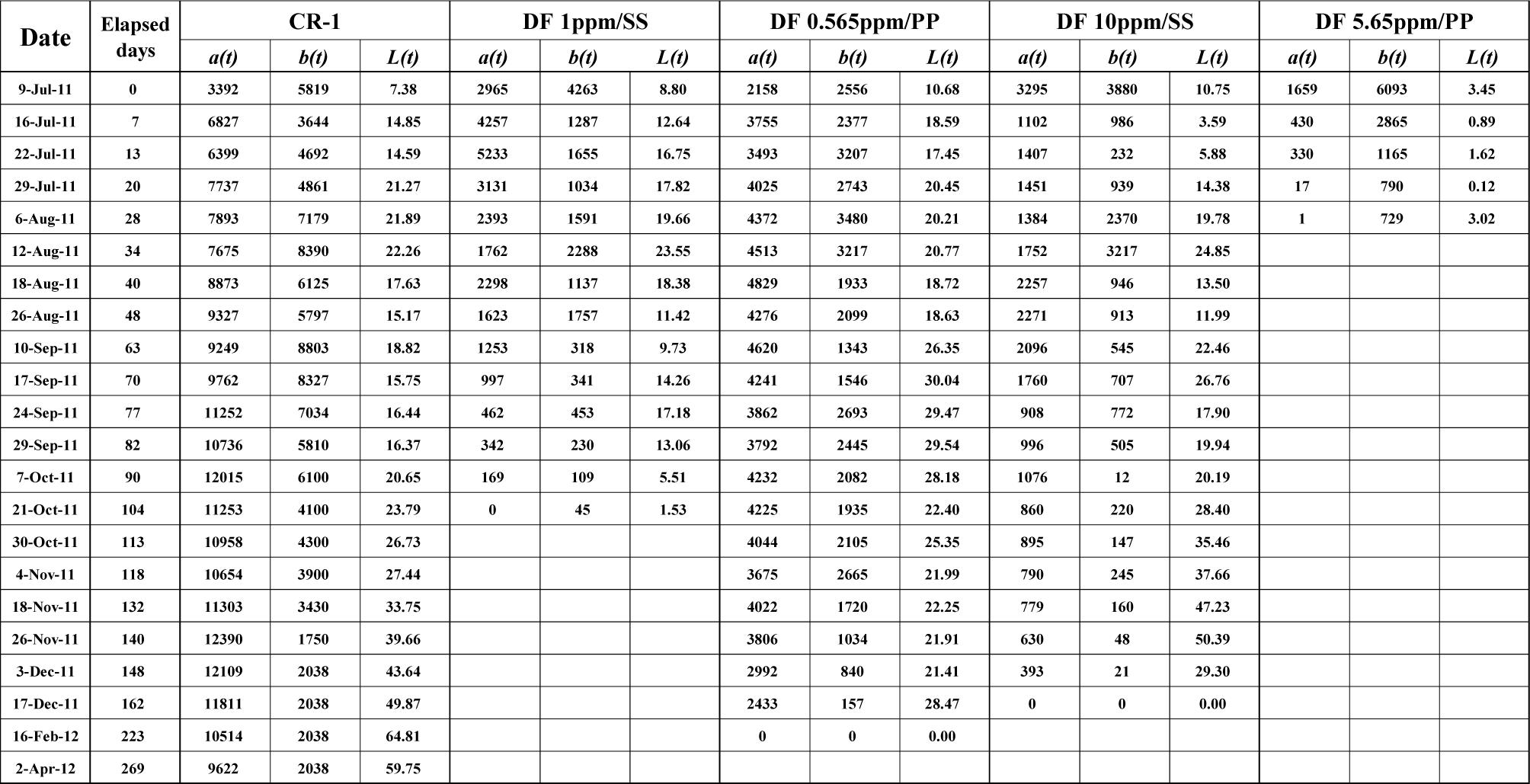
Apparent longevity of a honeybee colony in the 2011/2012 experiment at the eclosion rate of 0.9. This table shows the data source of the apparent longevity in Figure S7, which were estimated using the field experimental results conducted in 2011 to 2012. *a(t)* and *b(t)* denote the number of adult bees and the number of capped brood obtained from the field experiment conducted from 2011 to 2012, respectively (T. Yamada et al., 2018a). *L(t)* is the apparent longevity estimated using *a(t)* and *b(t)* by the mathematical model previously proposed (*1*). “CR-1” denotes the control colony where mites are almost absent and a pesticide is not administered. “DF 1ppm/SS” and “DF 10ppm/SS” denote the colonies where the neonicotinoid dinotefuran of 1 ppm and 10 ppm are administered via sugar syrup, respectively. “DF 0.565ppm/PP” and “DF 5.65ppm/PP” denote the colonies where the neonicotinoid dinotefuran of 0.565 ppm and 5.65 are administered via pollen paste, respectively. **Note:** A pesticide-administration period is from the morning of July 9 in 2011 until a colony becomes extinct, but until the morning of December 3 when a colony survives to the winter (T. Yamada et al., 2018a).

| Date | Elapsed days | CR-1 |  |  | DF 1ppm/SS |  |  | DF 0.565ppm/PP |  |  | DF 10ppm/SS |  |  | DF 5.65ppm/PP |  |  |
| --- | --- | --- | --- | --- | --- | --- | --- | --- | --- | --- | --- | --- | --- | --- | --- | --- |
| | | $a(t)$ | $b(t)$ | $L(t)$ | $a(t)$ | $b(t)$ | $L(t)$ | $a(t)$ | $b(t)$ | $L(t)$ | $a(t)$ | $b(t)$ | $L(t)$ | $a(t)$ | $b(t)$ | $L(t)$ |
| 9-Jul-11 | 0 | 3392 | 5819 | 7.38 | 2965 | 4263 | 8.80 | 2158 | 2556 | 10.68 | 3295 | 3880 | 10.75 | 1659 | 6093 | 3.45 |
| 16-Jul-11 | 7 | 6827 | 3644 | 14.85 | 4257 | 1287 | 12.64 | 3755 | 2377 | 18.59 | 1102 | 986 | 3.59 | 430 | 2865 | 0.89 |
| 22-Jul-11 | 13 | 6399 | 4692 | 14.59 | 5233 | 1655 | 16.75 | 3493 | 3207 | 17.45 | 1407 | 232 | 5.88 | 330 | 1165 | 1.62 |
| 29-Jul-11 | 20 | 7737 | 4861 | 21.27 | 3131 | 1034 | 17.82 | 4025 | 2743 | 20.45 | 1451 | 939 | 14.38 | 17 | 790 | 0.12 |
| 6-Aug-11 | 28 | 7893 | 7179 | 21.89 | 2393 | 1591 | 19.66 | 4372 | 3480 | 20.21 | 1384 | 2370 | 19.78 | 1 | 729 | 3.02 |
| 12-Aug-11 | 34 | 7675 | 8390 | 22.26 | 1762 | 2288 | 23.55 | 4513 | 3217 | 20.77 | 1752 | 3217 | 24.85 |  |  |  |
| 18-Aug-11 | 40 | 8873 | 6125 | 17.63 | 2298 | 1137 | 18.38 | 4829 | 1933 | 18.72 | 2257 | 946 | 13.50 |  |  |  |
| 26-Aug-11 | 48 | 9327 | 5797 | 15.17 | 1623 | 1757 | 11.42 | 4276 | 2099 | 18.63 | 2271 | 913 | 11.99 |  |  |  |
| 10-Sep-11 | 63 | 9249 | 8803 | 18.82 | 1253 | 318 | 9.73 | 4620 | 1343 | 26.35 | 2096 | 545 | 22.46 |  |  |  |
| 17-Sep-11 | 70 | 9762 | 8327 | 15.75 | 997 | 341 | 14.26 | 4241 | 1546 | 30.04 | 1760 | 707 | 26.76 |  |  |  |
| 24-Sep-11 | 77 | 11252 | 7034 | 16.44 | 462 | 453 | 17.18 | 3862 | 2693 | 29.47 | 908 | 772 | 17.90 |  |  |  |
| 29-Sep-11 | 82 | 10736 | 5810 | 16.37 | 342 | 230 | 13.06 | 3792 | 2445 | 29.54 | 996 | 505 | 19.94 |  |  |  |
| 7-Oct-11 | 90 | 12015 | 6100 | 20.65 | 169 | 109 | 5.51 | 4232 | 2082 | 28.18 | 1076 | 12 | 20.19 |  |  |  |
| 21-Oct-11 | 104 | 11253 | 4100 | 23.79 | 0 | 45 | 1.53 | 4225 | 1935 | 22.40 | 860 | 220 | 28.40 |  |  |  |
| 30-Oct-11 | 113 | 10958 | 4300 | 26.73 |  |  |  | 4044 | 2105 | 25.35 | 895 | 147 | 35.46 |  |  |  |
| 4-Nov-11 | 118 | 10654 | 3900 | 27.44 |  |  |  | 3675 | 2665 | 21.99 | 790 | 245 | 37.66 |  |  |  |
| 18-Nov-11 | 132 | 11303 | 3430 | 33.75 |  |  |  | 4022 | 1720 | 22.25 | 779 | 160 | 47.23 |  |  |  |
| 26-Nov-11 | 140 | 12390 | 1750 | 39.66 |  |  |  | 3806 | 1034 | 21.91 | 630 | 48 | 50.39 |  |  |  |
| 3-Dec-11 | 148 | 12109 | 2038 | 43.64 |  |  |  | 2992 | 840 | 21.41 | 393 | 21 | 29.30 |  |  |  |
| 17-Dec-11 | 162 | 11811 | 2038 | 49.87 |  |  |  | 2433 | 157 | 28.47 | 0 | 0 | 0.00 |  |  |  |
| 16-Feb-12 | 223 | 10514 | 2038 | 64.81 |  |  |  | 0 | 0 | 0.00 |  |  |  |  |  |  |
| 2-Apr-12 | 269 | 9622 | 2038 | 59.75 |  |  |  |  |  |  |  |  |  |  |  |  |

**Supplementary Table S4.** Apparent longevity of a honeybee colony in the 2012/2013 experiment at the eclosion rate of 0.9. This table shows the data source of the apparent longevity in Figure S8, which were estimated using the field experimental results conducted in 2012 to 2013. *a(t)* and *b(t)* denote the number of adult bees and the number of capped brood obtained from the field experiment conducted from 2012 to 2013, respectively (T. Yamada et al., 2018b). *L(t)* is the apparent longevity estimated using *a(t)* and *b(t)* by the mathematical model previously proposed (*1*). “CR-1” and “CR-2” denote the control colonies where a pesticide is not administered, respectively. “DF 2ppm/SS” denotes the colonies where the neonicotinoid dinotefuran of 2 ppm is administered via sugar syrup. “FT 10ppm/SS” denotes the colony where the organophosphate fenitrothion of 10 ppm is administered via sugar syrup. Mites did not little exist in all colonies. **Note:** A pesticide-administration period is from the morning of July 21 in 2012 until the morning of August 16 (T. Yamada et al., 2018b).

| Date | Elapsed days | CR-1 |  |  | CR-2 |  |  | DF 2ppm/SS |  |  | FT 10ppm/SS |  |  |
| --- | --- | --- | --- | --- | --- | --- | --- | --- | --- | --- | --- | --- | --- |
| | | $a(t)$ | $b(t)$ | $L(t)$ | $a(t)$ | $b(t)$ | $L(t)$ | $a(t)$ | $b(t)$ | $L(t)$ | $a(t)$ | $b(t)$ | $L(t)$ |
| 28-Jun-12 | 0 | 7136 | 4746 | 19.03 | 5832 | 5094 | 14.49 | 7119 | 5679 | 15.86 | 5690 | 4012 | 17.95 |
| 8-Jul-12 | 10 | 9621 | 5806 | 25.65 | 8917 | 4710 | 22.15 | 8877 | 6446 | 19.78 | 7157 | 5286 | 22.58 |
| 15-Jul-12 | 17 | 8695 | 10215 | 21.94 | 8265 | 11143 | 21.07 | 6878 | 13267 | 14.61 | 7565 | 9619 | 22.06 |
| 21-Jul-12 | 23 | 9647 | 10254 | 21.53 | 9665 | 11301 | 23.06 | 9173 | 9442 | 16.91 | 8943 | 8732 | 22.56 |
| 22-Jul-12 | 24 | 10136 | 10210 | 21.12 | 9558 | 10967 | 20.76 | 4885 | 8834 | 6.11 | 7750 | 8694 | 16.76 |
| 27-Jul-12 | 29 | 10633 | 10617 | 14.51 | 10770 | 10329 | 13.24 | 1434 | 4548 | 1.03 | 8721 | 6563 | 11.57 |
| 28-Jul-12 | 30 | 10391 | 10858 | 13.79 | 10901 | 10581 | 13.15 | 1313 | 3891 | 1.90 | 7786 | 6389 | 9.52 |
| 3-Aug-12 | 36 | 12083 | 10000 | 15.41 | 11939 | 10025 | 13.52 | 1049 | 1131 | 5.15 | 8289 | 3390 | 12.31 |
| 4-Aug-12 | 37 | 12389 | 9687 | 14.88 | 12041 | 10269 | 13.70 | 771 | 840 | 1.22 | 7559 | 2901 | 11.96 |
| 8-Aug-12 | 41 | 14065 | 7154 | 16.23 | 12978 | 8472 | 14.86 | 33 | 208 | 0.05 | 6625 | 1352 | 12.62 |
| 16-Aug-12 | 49 | 13371 | 6111 | 18.39 | 12207 | 5977 | 15.60 | 0 | 0 | 0.00 | 5961 | 607 | 18.86 |
| 25-Aug-12 | 58 | 11961 | 6014 | 19.88 | 10997 | 6684 | 18.37 |  |  |  | 4467 | 918 | 22.64 |
| 6-Sep-12 | 70 | 11165 | 8783 | 23.00 | 11582 | 8126 | 21.68 |  |  |  | 3534 | 3406 | 31.86 |
| 15-Sep-12 | 79 | 11980 | 5531 | 21.31 | 11825 | 7135 | 20.47 |  |  |  | 4576 | 4187 | 38.45 |
| 21-Sep-12 | 85 | 12166 | 6086 | 21.00 | 11025 | 9202 | 18.04 |  |  |  | 5859 | 4119 | 36.60 |
| 5-Oct-12 | 99 | 10715 | 7615 | 21.14 | 10510 | 5679 | 16.13 |  |  |  | 6593 | 4648 | 20.80 |
| 19-Oct-12 | 113 | 11726 | 7280 | 19.59 | 10038 | 6628 | 19.03 |  |  |  | 7326 | 4713 | 20.95 |
| 25-Nov-12 | 150 | 13255 | 36 | 47.53 | 12477 | 2937 | 37.74 |  |  |  | 7755 | 10 | 45.86 |
| 13-Dec-12 | 168 | 12858 | 0 | 64.81 | 13316 | 305 | 51.75 |  |  |  | 7080 | 0 | 62.00 |
| 1-Feb-13 | 218 | 9306 | 0 | 108.91 | 9421 | 0 | 93.76 |  |  |  | 5652 | 0 | 108.11 |
| 1-Mar-13 | 246 | 7464 | 17 | 133.75 | 8426 | 0 | 118.44 |  |  |  | 5957 | 26 | 136.85 |
| 9-Mar-13 | 254 | 7512 | 497 | 141.82 | 8017 | 660 | 124.67 |  |  |  | 6358 | 619 | 145.90 |
| 17-Mar-13 | 262 | 6862 | 1691 | 148.28 | 7372 | 2419 | 128.57 |  |  |  | 6152 | 2329 | 152.54 |
| 23-Mar-13 | 268 | 7312 | 2431 | 154.08 | 7416 | 3624 | 131.34 |  |  |  | 6470 | 3217 | 157.36 |
| 29-Mar-13 | 274 | 7720 | 4097 | 159.04 | 8018 | 5165 | 133.66 |  |  |  | 7143 | 4994 | 161.37 |
| 7-Apr-13 | 283 | 9518 | 7326 | 166.80 | 9833 | 8414 | 136.13 |  |  |  | 8797 | 7053 | 166.52 |
| 13-Apr-13 | 289 | 12523 | 10166 | 172.59 | 12594 | 11211 | 138.63 |  |  |  | 12038 | 8670 | 172.62 |
| 19-Apr-13 | 295 | 15677 | 11324 | 177.64 | 16221 | 12975 | 141.88 |  |  |  | 14275 | 10320 | 175.61 |
| 26-Apr-13 | 302 | 20574 | 9725 | 181.88 | 20412 | 14287 | 38.93 |  |  |  | 17132 | 10803 | 37.89 |
| 3-May-13 | 309 | 23935 | 5808 | 185.32 | 24100 | 14521 | 30.47 |  |  |  | 20413 | 9631 | 33.22 |
| 10-May-13 | 316 | 23629 | 4551 | 34.93 | 27670 | 13380 | 28.37 |  |  |  | 16477 | 9020 | 20.60 |
| 17-May-13 | 323 | 16210 | 6036 | 26.01 | 30325 | 12864 | 28.76 |  |  |  | 11103 | 8732 | 14.91 |
| 26-May-13 | 332 | 9981 | 9778 | 22.57 | 17904 | 10157 | 17.50 |  |  |  | 8817 | 6867 | 12.86 |
| 2-Jun-13 | 339 | 9289 | 7396 | 16.76 | 26814 | 3454 | 27.14 |  |  |  | 5763 | 5069 | 9.89 |
| 14-Jun-13 | 351 | 6135 | 4875 | 11.18 | 16284 | 1600 | 26.34 |  |  |  | 3936 | 2720 | 10.49 |
| 21-Jun-13 | 358 | 5590 | 3089 | 14.36 | 11501 | 7725 | 27.64 |  |  |  | 3058 | 1706 | 11.92 |
| 28-Jun-13 | 365 | 4210 | 1522 | 12.24 | 10454 | 10165 | 31.03 |  |  |  | 2336 | 1477 | 12.21 |
| 8-Jul-13 | 375 | 1962 | 978 | 13.31 | 11162 | 7534 | 21.93 |  |  |  | 1839 | 796 | 14.87 |
| 26-Jul-13 | 393 |  |  |  | 4817 | 4498 | 10.72 |  |  |  |  |  |  |

**Supplementary Table S5.** Apparent longevity of a honeybee colony in the 2013/2014 experiment at the eclosion rate of 0.9. This table shows the data source of the apparent longevity in Figure S9, which were estimated using the field experimental results conducted in 2013 to 2014. *a(t)* and *b(t)* denote the number of adult bees and the number of capped brood obtained from the field experiment conducted from 2012 to 2013, respectively (T. Yamada et al., 2018c). *L(t)* is the apparent longevity estimated using *a(t)* and *b(t)* by the mathematical model previously proposed (*1*). “CR-1” and “CR-2” denotes the mite-free control colonies where a pesticide is not administered. “DF 0.2ppm/SS” denotes the mite-free colony where the neonicotinoid dinotefuran of 0.2 ppm is administered via sugar syrup. “CN 0.0.08ppm/SS” denotes the mite-free colony where the neonicotinoid clothianidin of 0.08 ppm is administered via sugar syrup. “FT 1ppm/SS” denotes the mite-free colony where the organophosphate fenitrothion of 1 ppm is administered via sugar syrup. “MT 1ppm/SS” denotes the mite-free colony where the organophosphate malathion of 1 ppm is administered via sugar syrup. **Note:** A pesticide-administration period is from the morning of September 5 in 2013 until December 1 (T. Yamada et al., 2018c).

| Date | Elapsed days | CR-1 |  |  | DF 0.2 ppm/SS |  |  | FT 1 ppm/SS |  |  | MT 1 ppm/SS |  |  | CN 0.08 ppm/SS |  |  | CR-2 |  |  |
| --- | --- | --- | --- | --- | --- | --- | --- | --- | --- | --- | --- | --- | --- | --- | --- | --- | --- | --- | --- |
| | | $a(t)$ | $b(t)$ | $L(t)$ | $a(t)$ | $b(t)$ | $L(t)$ | $a(t)$ | $b(t)$ | $L(t)$ | $a(t)$ | $b(t)$ | $L(t)$ | $a(t)$ | $b(t)$ | $L(t)$ | $a(t)$ | $b(t)$ | $L(t)$ |
| 13-Aug-13 | 0 | 7579 | 4167 | 23.01 | 5447 | 4605 | 14.97 | 7360 | 5094 | 18.28 | 5500 | 4616 | 15.08 | 5628 | 5022 | 14.18 | 5953 | 4683 | 16.09 |
| 24-Aug-13 | 11 | 7000 | 3363 | 21.26 | 7556 | 323 | 20.76 | 9290 | 3758 | 23.08 | 6803 | 2414 | 18.65 | 6915 | 2208 | 17.43 | 6041 | 2146 | 16.32 |
| 1-Sep-13 | 19 | 7324 | 1825 | 23.77 | 6529 | 1219 | 25.07 | 10402 | 2671 | 27.90 | 8177 | 1054 | 26.11 | 7790 | 1025 | 23.95 | 5565 | 558 | 19.22 |
| 5-Sep-13 | 23 | 7487 | 2261 | 25.14 | 6017 | 2214 | 27.74 | 10296 | 3232 | 28.81 | 7676 | 1396 | 26.84 | 7293 | 1506 | 25.17 | 5549 | 578 | 21.56 |
| 15-Sep-13 | 33 | 6415 | 5462 | 27.73 | 5072 | 1743 | 30.73 | 5795 | 6040 | 22.27 | 5716 | 4992 | 29.32 | 5574 | 1825 | 28.67 | 4027 | 2665 | 26.86 |
| 21-Sep-13 | 39 | 6697 | 4683 | 27.08 | 5154 | 90 | 34.44 | 7431 | 3015 | 24.86 | 5698 | 3943 | 29.32 | 4001 | 843 | 28.20 | 3829 | 3198 | 29.13 |
| 27-Sep-13 | 45 | 5711 | 2538 | 13.68 | 3317 | 0 | 33.45 | 6920 | 1655 | 16.68 | 5617 | 2456 | 22.93 | 3885 | 208 | 30.74 | 3889 | 2882 | 29.88 |
| 4-Oct-13 | 52 | 5048 | 649 | 14.39 | 3145 | 0 | 41.41 | 5124 | 1456 | 17.33 | 4445 | 984 | 14.47 | 1882 | 160 | 19.35 | 3824 | 2142 | 16.20 |
| 13-Oct-13 | 61 | 4210 | 151 | 20.18 | 2029 | 0 | 31.82 | 4478 | 11 | 22.14 | 3459 | 0 | 18.90 | 1762 | 7 | 27.41 | 4322 | 113 | 20.63 |
| 27-Oct-13 | 75 | 3152 | 314 | 31.34 | 1325 | 0 | 41.98 | 3147 | 0 | 33.36 | 2328 | 0 | 30.19 | 1449 | 0 | 40.19 | 3371 | 0 | 29.89 |
| 15-Nov-13 | 94 | 2714 | 50 | 48.37 | 1146 | 0 | 60.17 | 2554 | 0 | 51.13 | 2050 | 0 | 47.95 | 1143 | 0 | 56.81 | 3181 | 0 | 48.19 |
| 1-Dec-13 | 110 | 2189 | 3 | 62.34 | 795 | 0 | 74.23 | 2213 | 0 | 66.43 | 1745 | 0 | 62.35 | 867 | 0 | 70.46 | 2904 | 0 | 63.18 |
| 5-Jan-14 | 145 | 1100 | 0 | 92.61 | 0 | 0 | 0.00 | 1714 | 0 | 100.39 | 916 | 0 | 92.97 | 0 | 0 | 0.00 | 2218 | 0 | 95.60 |
| 7-Feb-14 | 178 | 0 | 314 | 3.81 |  |  |  | 878 | 0 | 124.23 | 0 | 0 | 0.00 |  |  |  | 1799 | 0 | 127.15 |
| 28-Feb-14 | 199 | 0 |  | 0.00 |  |  |  | 0 | 0 | 0.00 |  |  |  |  |  |  | 1594 | 0 | 147.03 |

**Supplementary Table S6.** Apparent longevity, *L(t)*, of a honeybee colony in the 2018 experiment at the eclosion rate of 0.9. This table shows the data source of the apparent longevity in Figure S10, which were estimated using the field experimental results conducted in 2018. *a(t)* and *b(t)* denote the number of adult bees and the number of capped brood obtained from the field experiment conducted from 2012 to 2013, respectively (T. Yamada, 2020). *L(t)* is the apparent longevity estimated using *a(t)* and *b(t)* by the mathematical model previously proposed (*1*). “CR-1”, “CR-2” and “CR-3” denote the mite-infested control colonies where a pesticide is not administered. “DF-1 0.4ppm/SS”, “DF-2 0.4ppm/SS” and “DF-3 0.4ppm/SS” denote the mite-infested colonies where the neonicotinoid dinotefuran of 0.4 ppm is administered via pollen paste. Note: A pesticide-administration period is from the morning of July 1 in 2018 until the colony extinction (T. Yamada, 2020).

| Date | Elapsed days | CR-1 |  |  | CR-2 |  |  | CR-3 |  |  | DF-1 0.4ppm/PP |  |  | DF-2 0.4ppm/PP |  |  | DF-3 0.4ppm/PP |  |  |
| --- | --- | --- | --- | --- | --- | --- | --- | --- | --- | --- | --- | --- | --- | --- | --- | --- | --- | --- | --- |
| | | $a(t)$ | $b(t)$ | $L(t)$ | $a(t)$ | $b(t)$ | $L(t)$ | $a(t)$ | $b(t)$ | $L(t)$ | $a(t)$ | $b(t)$ | $L(t)$ | $a(t)$ | $b(t)$ | $L(t)$ | $a(t)$ | $b(t)$ | $L(t)$ |
| 19-Jun-18 | 0 | 8613 | 5341 | 20.41 | 8343 | 3065 | 34.45 | 6987 | 6861 | 12.89 | 9010 | 5891 | 19.36 | 9067 | 3241 | 35.40 | 7079 | 7726 | 678.30 |
| 1-Jul-18 | 12 | 10507 | 11056 | 24.90 | 8439 | 9401 | 34.84 | 9830 | 8621 | 18.13 | 10188 | 9341 | 21.89 | 8926 | 12549 | 34.85 | 10071 | 5374 | 471.80 |
| 16-Jul-18 | 27 | 13064 | 10657 | 15.13 | 12402 | 7447 | 22.11 | 12016 | 8060 | 18.56 | 12070 | 8360 | 17.64 | 13726 | 8755 | 14.75 | 10491 | 8032 | 705.20 |
| 28-Jul-18 | 39 | 12439 | 5688 | 14.77 | 11658 | 7640 | 18.81 | 10037 | 7006 | 15.71 | 11239 | 5443 | 16.80 | 11252 | 5769 | 15.88 | 10301 | 9052 | 794.80 |
| 12-Aug-18 | 54 | 6977 | 3817 | 15.81 | 9812 | 4619 | 17.50 | 6965 | 4078 | 13.83 | 8437 | 2777 | 18.96 | 8588 | 4030 | 18.13 | 9213 | 4840 | 425.00 |
| 28-Aug-18 | 70 | 6284 | 4125 | 20.01 | 9902 | 6629 | 22.86 | 6583 | 5277 | 19.15 | 7609 | 466 | 28.70 | 7558 | 5690 | 21.16 | 5662 | 4504 | 395.50 |
| 11-Sep-18 | 84 | 7369 | 4755 | 22.65 | 8589 | 5050 | 16.87 | 8081 | 6262 | 19.30 | 4716 | 395 | 34.80 | 8742 | 5928 | 19.92 | 4204 | 2978 | 261.50 |
| 23-Sep-18 | 96 | 6427 | 4201 | 17.58 | 9102 | 5636 | 20.71 | 6891 | 6487 | 13.93 | 2370 | 200 | 36.12 | 8046 | 6559 | 17.31 | 1158 | 1266 | 111.20 |
| 6-Oct-18 | 109 | 3680 | 1490 | 11.73 | 8593 | 5982 | 19.96 | 6150 | 5015 | 12.22 | 964 | 105 | 35.94 | 7599 | 6091 | 14.92 | 41 | 1228 | 107.80 |
| 23-Oct-18 | 126 | 1187 | 658 | 12.87 | 6395 | 4200 | 15.02 | 3461 | 2773 | 10.97 | 309 | 70 | 28.98 | 4366 | 3113 | 11.52 | 0 | 1234 | 108.30 |
| 4-Nov-18 | 138 | 596 | 440 | 11.46 | 3373 | 1747 | 10.16 | 1332 | 1043 | 6.08 | 204 | 43 | 30.13 | 1957 | 1646 | 7.96 | 0 | 0 | 0.10 |
| 18-Nov-18 | 152 | 207 | 411 | 6.09 | 383 | 913 | 3.73 | 254 | 607 | 3.92 | 112 | 39 | 25.76 | 191 | 1147 | 2.07 | 0 | 0 | 0.00 |
| 2-Dec-18 | 166 | 56 | 460 | 1.54 | 0 | 989 | 0.00 | 0 | 691 | 0.00 | 0 | 72 | 0.00 | 0 | 1291 | 0.00 | 0 | 0 | 0.00 |
| 16-Dec-18 | 180 | 0 | 463 | 0.00 | 0 | 0 | 0.00 | 0 | 0 | 0.00 | 0 | 0 | 0.00 | 0 | 0 | 0.00 | 0 | 0 | 0.00 |

**Supplementary Table S7.** Number of mite-damaged bees and mite-prevalence in 2018 and 2011/2012 experiments. 2018 (T. Yamada, 2020) and 2011/2012 (T. Yamada et al., 2018a) denote the years when the field experiments were conducted, respectively. CR-1, CR-2 and CR-3 denote the control colonies where no pesticide was administered in the 2018 experiment (T. Yamada, 2020), respectively. DF-1, DF-2 and DF-3 denote the colonies where dinotefuran of 0.4 ppm was administered through pollen paste in the 2018 experiment (T. Yamada, 2020), respectively. DF denotes the colony where dinotefuran of 0.565 ppm was administered through pollen paste in 2011/2012 experiment (T. Yamada et al., 2018a). Mites and Adults denote the number of mite-damaged bees and the number of adult bees, respectively. Mite-prev. denotes the mite-prevalence which is given by dividing the number of mite-damaged bees (Mites) into the number of adult bees (Adults). Incidentally, the numbers of mite-damaged bees (Mites) and adult bees (Adults) and the mite-prevalence (Mite-prev.) in the 2018 experiment (T. Yamada, 2020) and the number of adult bees (Adults) in the 2011/2012 experiment (T. Yamada et al., 2018a) have already been reported. The number of mite-damaged bees (Mites) and the mite-prevalence (Mite-prev.) in the 2011/2012 experiment were newly measured in this work using the photograph images (T. Yamada et al., 2018a) taken in the observation dates selected at intervals of about one month.

|  | 2018 |  |  |  |  |  |  |  |  |  |  |  |  |  |  |  |  |  | 2011/2012 |  |  |
| --- | --- | --- | --- | --- | --- | --- | --- | --- | --- | --- | --- | --- | --- | --- | --- | --- | --- | --- | --- | --- | --- |
| Date of observation | CR-1 |  |  | CR-2 |  |  | CR-3 |  |  | DF-1 |  |  | DF-2 |  |  | DF-3 |  |  | DF |  |  |
|  | Mites | Adults | Mite-prev. | Mites | Adults | Mite-prev. | Mites | Adults | Mite-prev. | Mites | Adults | Mite-prev. | Mites | Adults | Mite-prev. | Mites | Adults | Mite-prev. | Mites | Adults | Mite-prev. |
| 19-Jun | 1103 | 8613 | 0.1281 | 1577 | 8343 | 0.1890 | 1237 | 6987 | 0.1770 | 1281 | 9010 | 0.1422 | 1795 | 9067 | 0.1980 | 1940 | 7079 | 0.2741 |  |  |  |
| 1-Jul | 1190 | 10507 | 0.1133 | 1482 | 8439 | 0.1756 | 1986 | 9830 | 0.2020 | 1739 | 10188 | 0.1707 | 2603 | 8926 | 0.2916 | 2457 | 10071 | 0.2440 |  |  |  |
| 9-Jul |  |  |  |  |  |  |  |  |  |  |  |  |  |  |  |  |  |  | 84 | 2158 | 0.0389 |
| 16-Jul | 2263 | 13064 | 0.1732 | 2255 | 12402 | 0.1818 | 2670 | 12016 | 0.2222 | 2380 | 12070 | 0.1972 | 3532 | 13726 | 0.2573 | 3488 | 10491 | 0.3325 |  |  |  |
| 28-Jul | 3475 | 12439 | 0.2794 | 2401 | 11658 | 0.2060 | 3142 | 10037 | 0.3130 | 3351 | 11239 | 0.2982 | 3269 | 11252 | 0.2905 | 3881 | 10301 | 0.3768 |  |  |  |
| 6-Aug |  |  |  |  |  |  |  |  |  |  |  |  |  |  |  |  |  |  | 112 | 4372 | 0.0256 |
| 12-Aug | 1876 | 6977 | 0.2689 | 2620 | 9812 | 0.2670 | 2523 | 6965 | 0.3622 | 2689 | 8437 | 0.3187 | 3519 | 8588 | 0.4098 | 3819 | 9213 | 0.4145 |  |  |  |
| 28-Aug | 2636 | 6284 | 0.4195 | 3313 | 9902 | 0.3346 | 2442 | 6583 | 0.3710 | 2643 | 7609 | 0.3474 | 4008 | 7558 | 0.5303 | 2378 | 5662 | 0.4200 |  |  |  |
| 10-Sep |  |  |  |  |  |  |  |  |  |  |  |  |  |  |  |  |  |  | 90 | 4620 | 0.0195 |
| 11-Sep | 3507 | 7369 | 0.4759 | 4125 | 8589 | 0.4803 | 4438 | 8081 | 0.5492 | 3229 | 4716 | 0.6847 | 5988 | 8742 | 0.6850 | 3072 | 4204 | 0.7307 |  |  |  |
| 23-Sep | 4230 | 6427 | 0.6582 | 6531 | 9102 | 0.7175 | 5793 | 6891 | 0.8407 | 1985 | 2370 | 0.8376 | 6523 | 8046 | 0.8107 | 1002 | 1158 | 0.8653 |  |  |  |
| 6-Oct | 2903 | 3680 | 0.7889 | 7227 | 8593 | 0.8410 | 5408 | 6150 | 0.8793 | 815 | 964 | 0.8454 | 6919 | 7599 | 0.9105 | 39 | 41 | 0.9512 |  |  |  |
| 7-Oct |  |  |  |  |  |  |  |  |  |  |  |  |  |  |  |  |  |  | 104 | 4232 | 0.0246 |
| 23-Oct | 943 | 1187 | 0.7944 | 5575 | 6395 | 0.8718 | 2654 | 3461 | 0.7668 | 247 | 309 | 0.7994 | 3502 | 4366 | 0.8021 | 1 | 0 | 1.0000 |  |  |  |
| 4-Nov | 292 | 596 | 0.4899 | 2989 | 3373 | 0.8862 | 1240 | 1332 | 0.9309 | 198 | 204 | 0.9706 | 1766 | 1957 | 0.9024 | 0 | 0 |  | 111 | 3675 | 0.0302 |
| 18-Nov | 118 | 207 | 0.5700 | 363 | 383 | 0.9478 | 245 | 254 | 0.9646 | 100 | 112 | 0.8929 | 172 | 191 | 0.9005 | 0 | 0 |  |  |  |  |
| 2-Dec | 52 | 56 | 0.9286 | 1 | 0 | 1.0000 | 1 | 0 | 1.0000 | 3 | 0 | 1.0000 | 1 | 0 | 1.0000 | 0 | 0 |  |  |  |  |
| 3-Dec |  |  |  |  |  |  |  |  |  |  |  |  |  |  |  |  |  |  | 144 | 2992 | 0.0481 |
| 16-Dec | 0 | 0 | 1.0000 |  |  |  |  |  |  |  |  |  |  |  |  | 0 | 0 |  |  |  |  |

